# SAFB2 enables the processing of suboptimal stem-loop structures in clustered primary miRNA transcripts

**DOI:** 10.1101/858647

**Authors:** Katharina Hutter, Michael Lohmüller, Almina Jukic, Felix Eichin, Seymen Avci, Verena Labi, Simon M. Hoser, Alexander Hüttenhofer, Andreas Villunger, Sebastian Herzog

## Abstract

MicroRNAs (miRNAs) are small noncoding RNAs that post-transcriptionally silence most protein-coding genes in mammals. They are generated from primary transcripts containing single or multiple clustered stem-loop structures that are thought to be recognized and cleaved by the DGCR8/DROSHA Microprocessor complex as independent units. Contrasting this view, we here report an unexpected mode of processing of a bicistronic cluster of the miR-15 family, miR-15a-16-1. We find that the primary miR-15a stem-loop is a poor Microprocessor substrate and is consequently not processed on its own, but that the presence of the neighboring primary miR-16-1 stem-loop on the same transcript can compensate for this deficiency in *cis*. Using a CRISPR/Cas9 screen, we identify SAFB2 (scaffold attachment factor B2) as an essential co-factor in this miR-16-1-assisted pri-miR-15 cleavage, and describe SAFB2 as a novel accessory protein of DROSHA. Notably, SAFB2-mediated cluster assistance expands to other clustered pri-miRNAs including miR-15b, miR-92a and miR-181b, indicating a general mechanism. Together, our study reveals an unrecognized function of SAFB2 in miRNA processing and suggests a scenario in which SAFB2 enables the binding and processing of suboptimal DGCR8/DROSHA substrates in clustered primary miRNA transcripts.

**Highlights:**

- the primary miR-15a stem-loop structure *per se* is a poor Microprocessor substrate
- cleavage of pri-miR-15a requires the processing of an additional miRNA stem-loop on the same RNA
- sequential pri-miRNA processing or “cluster assistance” is mediated by SAFB proteins
- SAFB2 associates with the Microprocessor

## Introduction

MicroRNAs (miRNAs) are 20-25 nucleotides (nt) RNAs that are predicted to post-transcriptionally silence expression of >60% of all protein coding genes in mammals (Friedman et al., 2008). In canonical miRNA biogenesis, transcription of miRNA genes by RNA polymerase II gives rise to long primary transcripts (pri-miRNAs) that contain one or several distinct stem-loop structures in which the mature miRNAs are embedded. The initial cleavage of the pri-miRNA in the nucleus is mediated by the Microprocessor complex, consisting of the RNase III-type enzyme DROSHA and its co-factor DGCR8 (DiGeorge syndrome critical region 8 homolog) (Denli et al., 2004; Gregory et al., 2004; Han et al., 2004; Landthaler et al., 2004; Lee et al., 2003; Nguyen et al., 2015). The Microprocessor binds and cleaves the stem-loop near its base to release a precursor miRNA (pre-miRNA) with a characteristic two nt 3’ overhang. Further processing of the pre-miRNA is mediated by the RNase III-type endonuclease DICER, which generates an RNA duplex that is loaded Argonaute family proteins (AGO) (Grishok et al., 2001; Hutvágner et al., 2001; Ketting et al., 2001). Removal of the passenger strand give rise to the mature RISC (RNA-induced silencing complex), the effector complex of miRNA-mediated gene regulation (J. Liu et al., 2004; Schwarz et al., 2003). The remaining guide strand directs the RISC to its target sites in mRNAs, ultimately resulting in gene repression by translational inhibition and/or mRNA degradation (Djuranovic et al., 2012; Friedman et al., 2008; Guo et al., 2010; Hafner et al., 2010).

As the initial step in miRNA biogenesis, the recognition of distinct pri-miRNA stem-loop structures and their subsequent cleavage by the Microprocessor needs to be tightly controlled. However, it remains unclear how the Microprocessor can distinguish between authentic primary miRNA stem-loops and the thousands of miRNA-like RNA folds within the entire transcriptome (Lim et al., 2003). Elegant studies have defined a set of features that either appear to be a prerequisite for processing or that have been shown to enhance Microprocessor-mediated cleavage in a modular or additive fashion. Structurally, these include a narrow stem length of 35±1 nt, a single-stranded apical loop of >10 nt, and an unstructured flanking region (Fang and Bartel, 2015; Han et al., 2006; Zeng and Cullen, 2005; Zeng et al., 2005). With respect to sequence motifs, high-throughput analyses have identified a basal UG motif at the 5’ boundary of single-stranded RNA flanking the lower stem, an UGU/GUG motif in the apical loop and a CNNC motif 16 to 18 nt downstream of the DROSHA cleavage site (Auyeung et al., 2013). More recently, a so-called mismatched GHG motif in the lower stem - a region that otherwise favors extensive base-pairing - has been added to the list (Fang and Bartel, 2015).

Notably, not all human pri-miRNAs contain such motifs, suggesting that other factors may enable Microprocessor-mediated processing. Indeed, various accessory proteins have been reported to either bind directly to the pri-miRNA or to the Microprocessor itself, thereby assisting in pri-miRNA cleavage (Alarcón et al., 2015; Cheng et al., 2014; Fletcher et al., 2016; Nussbacher and Yeo, 2018; Treiber et al., 2017; Tu et al., 2015). Moreover, recent studies also reveal stem-loop structures as *cis*-regulatory elements in pri-miRNA processing. Here, the processing of certain pri-miRNAs within polycistronic clusters depends on the presence of a second miRNA stem-loop on the same primary transcript, suggesting that cleavage of the latter licenses Microprocessor-mediated processing of the former. These clusters include *Drosophila* miR-11-998, in which deletion of miR-11 prevents the processing of pri-miR-998, as well as the highly conserved miR-100-let7-125, where miR-125 expression is enhanced by the presence of additional miRNA genes on the same pri-miRNA (Truscott et al., 2016). Likewise, CRISPR/Cas9-mediated editing of miR-195 in the miR-497-195 cluster led to a significant decrease in the expression of miR-497a, suggesting a sequential and co-dependent processing scheme (Lataniotis et al., 2017). Finally, efficient processing of the clustered Epstein-Barr virus (EBV) miR-BHRF1-3 appears to be tightly linked to its neighboring miR-BHRF1-2 (Haar et al., 2016).

Currently, it remains unclear how this co-dependent processing is mediated on the molecular level, i.e. what are the structural and/or sequence features that interfere with miRNA processing when expressed without an assisting stem-loop, and how can co-expression of a miRNA overcome this deficit? Is the assistance by a second stem-loop solely mediated by the Microprocessor, or are other co-factors involved? Equally important, it remains to be determined whether this phenomenon is a common feature of clustered miRNA biogenesis or rather an exception.

Here, we report that disrupting the hairpin structure of miR-16-1 within the miR-15a-16-1 cluster, best known for its tumor-suppressing function in chronic lymphocytic leukemia (CLL), results in a complete loss of pri- to pre-miR-15a processing. We show that this defective processing can be explained by an unpaired region in the lower stem of pri-miR-15a, a structure that interferes with efficient recognition by the Microprocessor and thus defines pri-miR-15a as a poor substrate. Using a genome-wide CRISPR/Cas9 loss-of-function screen to decipher how pri-miR-16-1 enables the cleavage of the suboptimal pri-miR-15a structure, we identify SAFB2 (scaffold attachment factor B2) as a critical factor for this cluster-mediated assistance. Notably, deletion of Safb1 and 2 genes reduces not only miR-15a levels, but also affects other clustered miRNAs, among them miR-15b, miR-181b and miR-92a, all of which appear to also represent suboptimal Microprocessor substrates. On the molecular level, we demonstrate that SAFB2 binds to the N-terminal regulatory region of DROSHA, and that loss of this interaction correlates with a loss-of-function in SAFB2-mediated miRNA biogenesis. We therefore postulate a model in which SAFB2 allows binding of the Microprocessor and/or processing of suboptimal substrates in a subset of clustered pri-miRNA transcripts.

## Results

### DGCR8/DROSHA-mediated processing of pri-miR-15a depends on the presence of a second miRNA stem-loop on the same primary transcript

Using a GFP-based reporter system that allows the flow cytometric quantification of mature miRNA activity, we initially verified that exogenous expression of the full miR-15a-16-1 cluster in Ramos cells resulted in clear target repression by miR-15a and miR-16, respectively (Fig. 1A and B). However, while miR-16-1 alone also induced efficient reporter repression, expression of only miR-15a did not produce any effect (Fig. 1B), suggesting a miR-16-1-dependent processing of the miR-15a stem-loop. To test this, we either destabilized the miR-16-1 stem-loop by introducing 4 mismatch mutations within its stem region (miR-15a_16-1^mut4^) or completely disrupted all base-pairing in the stem by mutagenesis, thereby inhibiting its processing by the DGCR8/DROSHA complex (Fig. 1C). When transduced into cells expressing the respective reporter, both constructs indeed showed a complete loss of miR-15a function compared to the wt (wild-type) cluster (Fig. 1C). Northern blot analysis on cells expressing the respective constructs demonstrated that no mature miR-15a was produced in the absence of functional miR-16-1 (Fig. 1D). Supporting this, a fluorescent reporter for the *in vivo* quantification of pri-miRNA processing by DGCR8/DROSHA (Allegra and Mertens, 2011) indicated that the miR-15a stem-loop was not processed without miR-16-1 (Fig. 1E and S1A). These data demonstrate that the pri- to pre-miR-15a cleavage depends on the presence of miR-16-1 on the same primary RNA transcript.

**Fig. 1:**
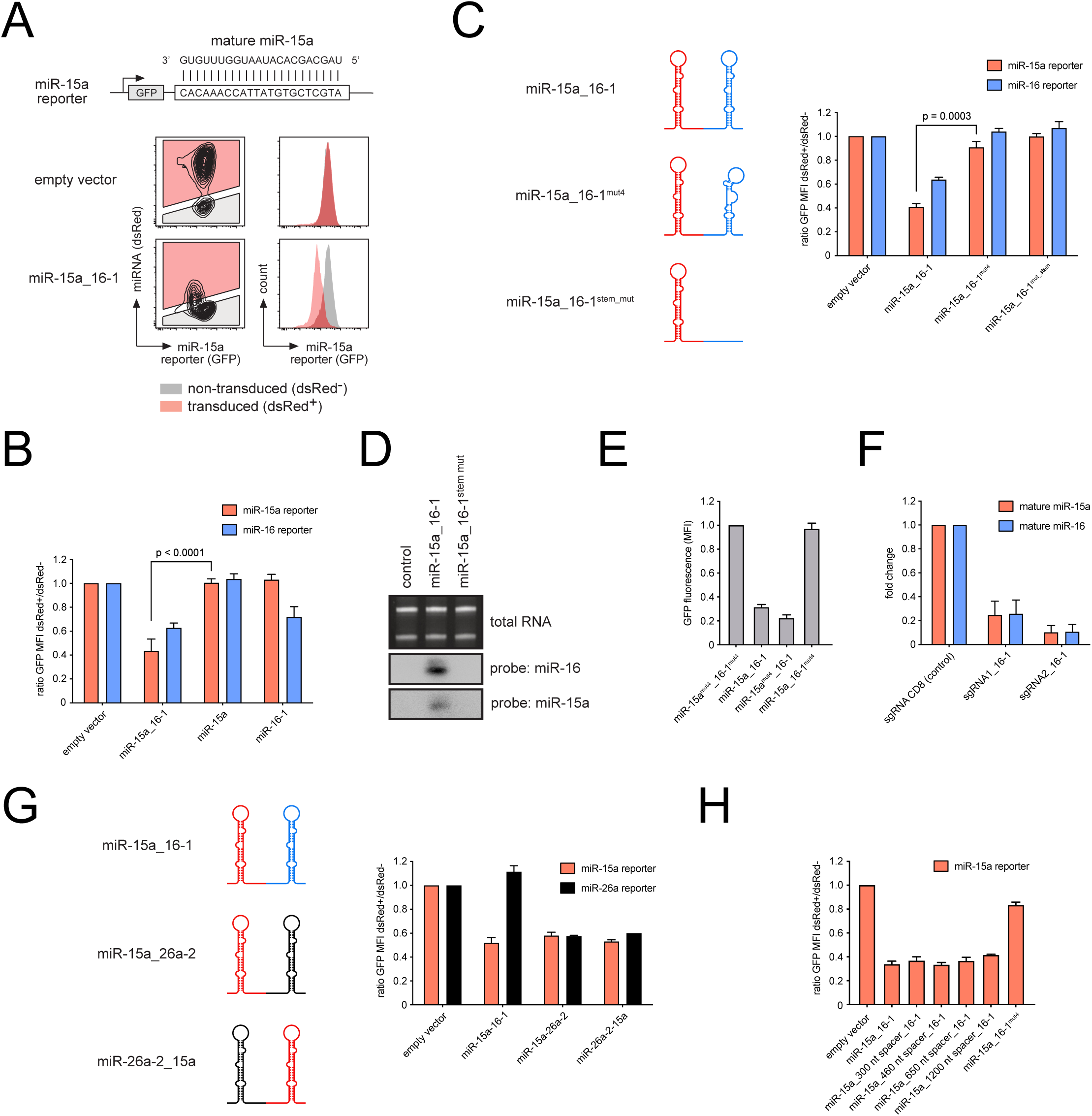
Primary miR-15a cleavage depends on the processing of a second miRNA stem-loop on the same transcript. **A.** Schematic illustration of the fluorescent reporter that quantifies mature miRNA activity. A sequence complementary to the miRNA of interest is cloned into the 3’-UTR of a cDNA encoding GFP. Co-expression of the reporter with the respective miRNA (dsRed as a marker) promotes GFP repression (lower panel), whereas a control vector has no effect (upper panel). **B.** MiR-15a is not active when expressed on its own. Ramos cells expressing miR-15a and miR-16 reporters were virally transduced with the indicated constructs and analyzed for GFP expression. **C**. MiRNA reporter assay as in B, using the illustrated wt and mutated miR-15a-16-1 clusters. **D**. Northern blot of 293T cells transfected with an empty control vector, a vector encoding the wt miR-15a-16-1 cluster or the cluster lacking the miR-16-1 stem-loop. Specific probes indicate expression of miR-15a and miR-16, respectively. Ribosomal RNA from the total RNA preparation is shown as a control. **E**. Primary miR-15 fails to be cleaved by the Microprocessor when expressed without a second stem-loop on the same RNA. Ramos cells were retrovirally transduced with constructs in which the depicted miR-15a-16-1 cluster variants were inserted in the 3’-UTR of a GFP cDNA (Fig. S1A). Individual bars depict the GFP MFI values (normalized to the miR-15a^mut4^_16-1^mut4^ control). **F**. Endogenous miR-15a requires the neighboring miR-16-1 for its biogenesis. Ramos clones in which the miR-16-2 locus was disrupted by CRISPR/Cas9 were lentivirally transduced with vectors encoding a control sgRNA or two individual sgRNAs targeting miR-16-1. After selection of transduced cells for one week, total RNA was isolated, reverse transcribed and subjected to qPCR for mature miR-15a and 16-1, respectively. **G**. Cluster assistance is independent of miR-16 function and orientation or distance of the assisting stem-loop. MiR-15a and miR-26a reporter Ramos cells were transduced with the depicted constructs and analyzed for GFP fluorescence. **H**. 293T cells stably expressing a miR-15a reporter were transfected with constructs encoding wt miR-15a_16-1 or mutants in which the individual miRNA stem-loops were spaced as indicated. GFP fluorescence was quantified by flow cytometry after 48 hrs. Data are represented as mean ± SD.

To investigate whether this assisted processing within a miRNA cluster, a process we termed “cluster assistance”, also applies to the endogenous miR-15a-16-1 gene, we employed a CRISPR/Cas9 approach. We first generated single cell clones lacking miR-16-2, which is expressed from the paralog cluster miR-15b-16-2, to ensure that all mature miR-16 in these cells originates from the miR-15a-16-1 loci. These cells were then transduced with a CRISPR/Cas9 construct targeting a control gene not involved in miRNA biogenesis, or with two individual sgRNAs (single guide RNA) targeting miR-16-1. As assessed by quantitative PCR (qPCR), loss of miR-16 in consequence of CRISPR/Cas9-mediated gene disruption by either sgRNA was accompanied by an equal loss of miR-15a (Fig. 1F), indicating that cluster assistance by miR-16-1 is indeed required for efficient biogenesis of endogenous miR-15a.

To define the basic principles of this unexpected regulation, we performed experiments with additional mutants of the miR-15a-16-1 cluster and found that the effect on miR-15a did not specifically depend on miR-16 function, since swapping the assisting miR-16-1 for the prototypic miR-26a-2 still allowed efficient cleavage of the primary miR-15a (Fig. 1G). Moreover, neither the relative localization of the assisting miRNA to miR-15a nor their distance within tested parameters affected processing of miR-15a (Fig. 1G and H), suggesting that any miRNA stem-loop that is located on the same primary transcript can license miR-15a cleavage in *cis*. However, such a stem-loop appears to require proper processing by DGCR8/DROSHA, as mutation of both the UG and CNNC motifs within the miR-16-1 context (Auyeung et al., 2013), two well-known motifs that enhance pri-miRNA processing in a modular fashion, completely abrogated miR-15a function (Fig. S1B). Thus, the mere presence of a stem-loop structure is not sufficient to induce cluster assistance.

Together, our data add pri-miR-15a to the growing list of clustered miRNAs in which processing of a second stem-loop structure on the same primary transcript licenses their own processing in *cis*. Since the underlying mechanism of this phenomenon is unclear, we wondered why miR-15a depends on processing of miR-16-1 and how exactly the presence of a second processable stem-loop facilitates miR-15a cleavage.

### An unpaired region within the lower stem prevents independent processing of pri-miR-15a

To address the former question, we first applied *in vivo* proximity biotin labeling (Ramanathan et al., 2018) to investigate whether the primary miR-15a *per se* fails to bind to the DGCR8/DROSHA complex, or whether it binds but is not properly cleaved. Using the miR-16-1 stem-loop as a positive and scrambled sequences as negative controls, these experiments demonstrated that in contrast to miR-16-1, miR-15a on its own does not attract the Microprocessor complex (Fig. 2A). This suggests that sequence features in either the pri-miR-15a context, i.e. in the region directly flanking the stem-loop, or in the stem-loop itself interfere with Microprocessor binding. To test the first possibility, we expressed the miR-26a-2 stem-loop flanked by miR-15a sequences and observed that miR-26a-2 was adequately cleaved independently of an assisting miR-16-1 (Fig.S2A), implicating that the miR-15a context is permissive for processing. To identify the features within the miR-15a stem-loop that hinder Microprocessor binding, we generated swapping mutants in which either the loop region or the lower stem of miR-15a were exchanged for the corresponding sequences of miR-26a-2 (Fig. 2B). While the loop mutant did not show any activity, replacing the lower stem of miR-15a for the corresponding sequences of miR-26a-2 resulted in strong reporter repression, suggesting that this region of miR-15a contains a feature that renders it a suboptimal or even defective Microprocessor substrate. Strikingly, compared to the prototypic miR-26a-2, the predicted secondary structure of miR-15a revealed an atypical extended region without base pairing in the lower stem (Fig. 2C), a structure that clearly opposes the experimentally defined requirements for primary miRNAs (Auyeung et al., 2013; Han et al., 2006; Lee et al., 2003).

**Figure 2:**
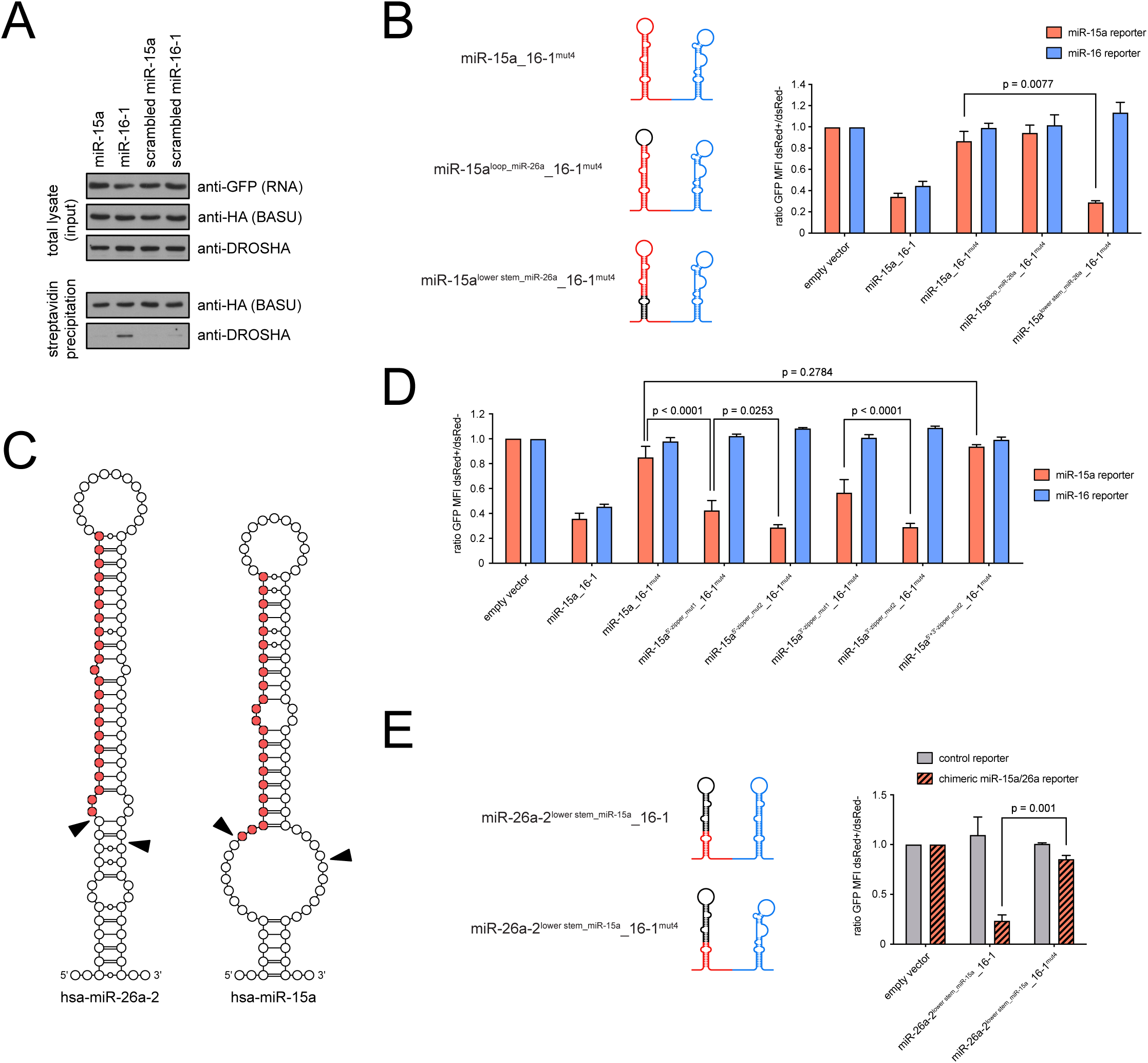
An unpaired region in its lower stem makes pri-miR-15a a poor substrate for the Microprocessor complex. **A**. Proximity labeling demonstrates recruitment of DROSHA to miR-16-1, but not to miR-15a. 293T cells transfected with the HA-tagged BASU biotin ligase and constructs encoding BoxB-flanked miR-15a and miR-16 or scrambled controls thereof were incubated with biotin for 4 hrs, lysed and subjected to a streptavidin pulldown. Western blots show the expression levels of the miRNA baits (GFP), the biotin ligase (HA) and endogenous DROSHA. HA-tagged BASU becomes auto-biotinylated and serves as a positive control for the pulldown. **B**. A feature in its lower stem renders pri-miR-15a a poor Microprocessor substrate. MiRNA reporter assay using chimeric miR-15a/miR-26a_16-1 contructs as depicted. Reporter repression was quantified by flow cytometry 48 hrs after transduction. **C**. Schematic illustration of the stem-loop portion of primary miR-26a-a and miR-15a. Red circles correspond to the respective mature miRNA. Microprocessor cleavage sites are marked by black triangles. **D**. Pairing mutagenesis of the miR-15a lower stem enables miR-16-1-independent processing of pri-miR-15a. Ramos cells were transduced with miR-15a_16-1 cluster mutants that increased the pairing in the lower stem (Fig. S2B) and were analyzed for reporter repression after 48 hrs. **E**. A chimeric construct that combines the miR-15a lower stem with the loop and upper stem of prototypic miR-26a-2 relies on cluster assistance for efficient miRNA processing. Ramos cells expressing a control reporter or a reporter against a chimeric miR-15a/26a were transduced with the depicted constructs (Fig. S2D) and analysed by flow cytometry. Data are represented as mean ± SD.

To test whether this unpaired region was indeed responsible for miR-15a’s inability to be processed by the DGCR8/DROSHA complex, we introduced either one or two pairing point mutations into the 5’ and the 3’ portion of the lower stem and expressed the respective clusters containing non-functional miR-16-1 in reporter cells (Fig. 2D and S2B). Remarkably, while the two single point mutations already had significant effects on miR-15a activity, introduction of two pairing mutations completely abolished the dependence of miR-15a on cluster assistance (Fig. 2D), i.e. mutated miR-15a in the absence of functional miR-16-1 was as efficient in repressing the reporter as was the wt cluster. However, a combination of all 4 pairing mutations, which were designed to mimic the unpaired situation as in the original miR-15a stem-loop, again abrogated miR-15a processing in the absence of miR-16-1 (Fig. 2D and S2B). This strongly indicates that the unpaired structure, rather than a sequence motif in this region, precludes Microprocessor binding.

These results made us wonder whether an unpaired region within the lower stem imposes the need for cluster assistance in general. However, when disrupting base pairing within the lower stem of miR-26a-2 by mutagenesis, such constructs were found to be inactive independent of an assisting miRNA on the same primary transcript (Fig. S2C). This suggests that the lower stem region of miR-15a contains features or adopts a structure that specifically enables its efficient processing only when expressed in the context of a miRNA cluster. Indeed, swapping the miR-15a lower stem onto miR-26a-2 generated a chimeric construct that showed little activity on its own, but that provoked strong reporter repression when expressed together with miR-16-1 (Fig. 2E and S2D).

Together, these experiments suggest that an unpaired region within the lower stem of miR-15a is on the one hand responsible for the defective DGCR8/DROSHA binding in the absence of an assisting miRNA, but on the other hand this particular region enables its processing in a clustered setup. This raised the question of how cluster assistance is mediated on the molecular level.

### Development of a genome-wide CRISPR/Cas9 screen to identify genes that mediate pri-miR-15a processing

Given that isolated pri-miR-15a failed to bind to DROSHA, we wondered whether the Microprocessor complex might be initially recruited via miR-16-1, serving as an entry point onto the primary transcript. Processing of pri-miR-16 might then allow DGCR8/DROSHA to relocate to the neighboring pri-miR-15a and might overcome its inability to independently recruit the enzymatic machinery. In this scenario, we furthermore hypothesized that the composition of the Microprocessor complex mediating processing of miR-15a differs from the one cleaving miR-16-1, i.e. that a co-factor alters the behavior of DGCR8 and/or DROSHA.

To pursue this hypothesis, we established an unbiased genome-wide CRISPR/Cas9 loss-of-function screen to identify genes specifically involved in miR-15a biogenesis (Fig. 3A). This screen was based on the combination of GFP- and dsRed-based miRNA reporters with a mutated miR-15a-16-1 cluster. In particular, we disrupted the miR-15a and miR-16-1 seed sequences by introduction of two point mutations in the 5’ strand and the corresponding positions on the 3’ strand, thereby avoiding miR-15a/16-1 target repression but preserving the folding of both stem-loop structures (Fig. S3A). This mutagenesis strategy served two purposes: First, exogenous expression of the wt miR-15a-16-1 cluster normally results in a profound cell-cycle arrest. Hence, the disruption of the seed sequences enabled us to express the cluster at high levels. Second, since the modified miRNAs were not expressed endogenously, we were able to optimize the dynamic range between “high miRNA activity” and “no miRNA activity”, which we considered as a critical parameter for a successful screen. Baf3 cells, a murine B lymphocyte progenitor cell line that showed the same cluster assistance phenotype as described for miR-15a above, were initially transduced with the two reporters for the mutated miR-15a (miR-15a^mut^, dsRed-based) and miR-16 (miR-16^mut^, GFP-based; Fig. 3A and data not shown). We then introduced the modified miR-15a^mut^-16-1^mut^ cluster into such GFP^+^dsRed^+^ reporter cells, inducing a repression of both fluorescent markers and thus generating a GFP^-^dsRed^-^ population that was FACS-sorted and hereafter is referred to as “screen cells” (Fig. 3A). Using these cells in a CRISPR/Cas9 loss-of-function screen, one would expect the targeted genes to fall into three different categories: The majority of genes is expected to have no function in general or specific miR-15a and/or miR-16-1 biogenesis, and thus should not induce derepression of the fluorescent reporters (Fig. 3B, upper panel). SgRNAs targeting the general miRNA biogenesis machinery, such as DGCR8 or DROSHA, or factors that are specifically involved in miR-16-1 biogenesis should derepress both reporters, since miR-15a is either directly affected or due to the loss of miR-16-1. The corresponding cells would therefore be expected to shift to the GFP^+^dsRed^+^ upper right quadrant (Fig. 3B, center panel). Most revealing for our question are the genes that are specific for miR-15a biogenesis, as such candidates would likely be involved in cluster assistance. If such a gene is disrupted, the respective population should become dsRed^+^, but remain GFP^-^ since the miR-16-1^mut^ reporter remains repressed (Fig. 3C, lower panel).

**Figure 3:**
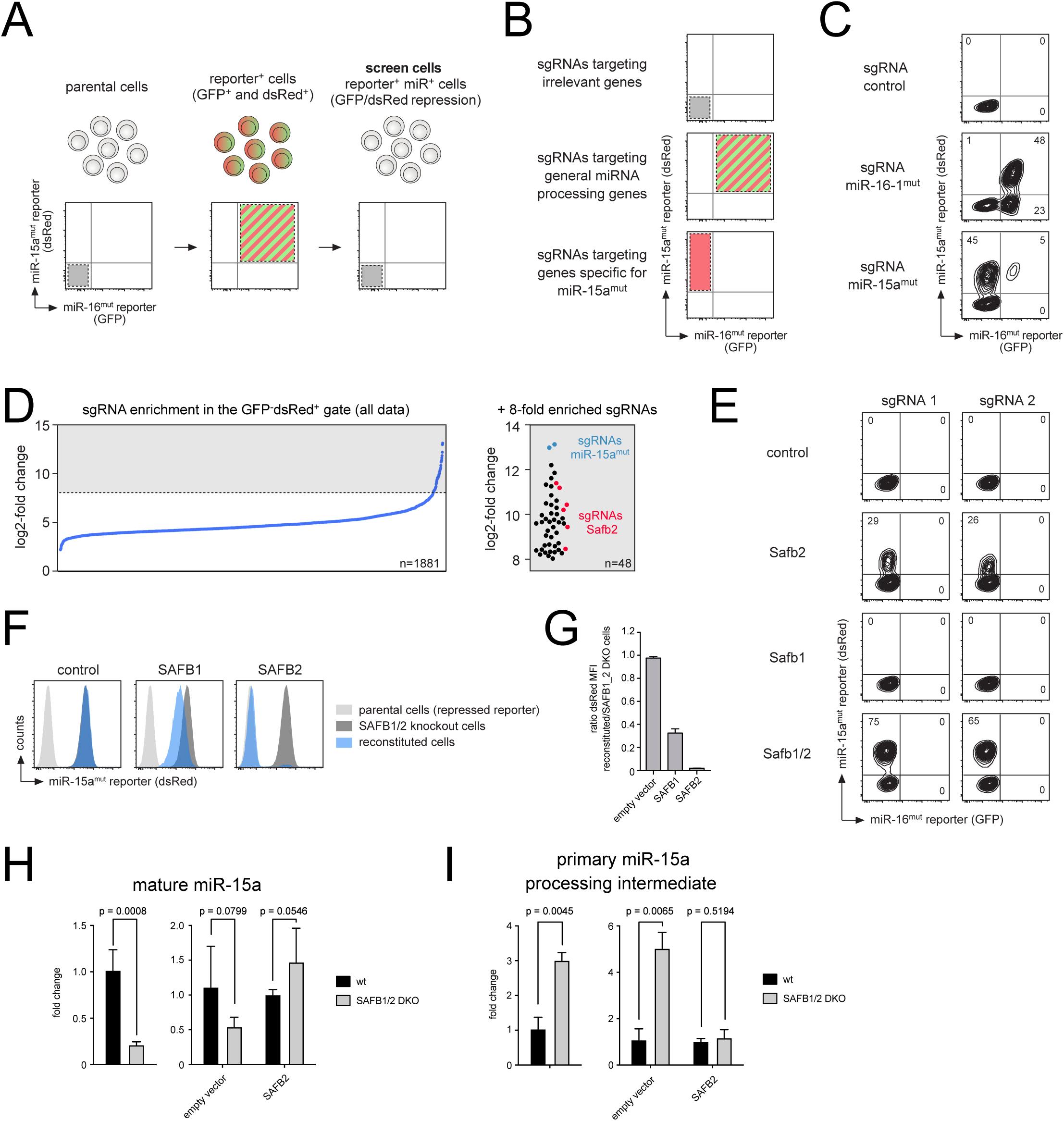
SAFB2 is required for efficient processing of primary miR-15a. **A**. Schematic illustration of the dual-fluorescence reporter cells used for the CRISPR/Cas9 screen. **B**. Expected cellular shifts upon CRISPR/Cas9-mediated disruption of general miRNA or miR-15a-specific regulatory genes. **C**. The cellular system can discriminate between general and specific miRNA regulators. Screen cells were lentivirally transduced with constructs encoding a control sgRNA or with sgRNAs targeting miR-16-1^mut^ or miR-15a^mut^. After one week, the fluorescence pattern was analyzed by flow cytometry. Numbers represent the percentage of cells within the respective quadrant. **D**. SgRNAs targeting the Safb2 gene are strongly enriched when repression of the miR-15a^mut^ reporter is lost. Enrichment of sgRNAs expression in the GFP^-^dsRed^+^ quadrant compared to the GFP^-^dsRed^-^ control for all data (left panel) and the top hits (> 8-fold enriched). Spiked-in sgRNAs targeting miR-15a^mut^ as a positive control are shown in blue, the six individual sgRNAs targeting Safb2 are depicted in red. **E**. Deletion of family members SAFB1 and SAFB2 validates their impact on miR-15a activity. Screen cells were lentivirally transduced with independent sgRNA constructs targeting control genes, Safb2 only, Safb1 only or both. After one week, reporter expression was measured by flow cytometry. **F**, **G**. Loss of miR-15a^mut^ reporter repression upon Safb1/2 disruption is restored by expression of full-length SAFB2. GFP^-^dsRed^+^ screen cells whose Safb1 and Safb2 genes had been disrupted were sorted and reconstituted with an empty vector control, human SAFB1 or human SAFB2. The calculated ratio of the dsRed MFI for reconstituted versus non-reconstituted cells of three individual experiments is provided as a bar graph. **H**, **I**. SAFB proteins are required for efficient pri- to pre-miRNA cleavage of endogenous miR-15a. Safb1/2 DKO and wt control cells were analyzed for mature miR-15a expression (H) or expression of a miR-15a-specific processing intermediate (I, Fig. S3E) by quantitative RT-PCR (left panels). Correspondingly, control cells and Safb1/2 DKO cells reconstituted with an empty vector or with a vector encoding a CRISPR-resistant variant of human SAFB2 for one week were subjected to RT-PCR analysis for the respective RNAs (H, I, right panels). Data are represented as mean ± SD.

To test our experimental setup, we transduced screen cells with constructs expressing Cas9 together with a control sgRNA and two sgRNAs targeting the mutated miR-16-1^mut^ and miR-15a^mut^ and analyzed the fluorescence shifts by flow cytometry (Fig. 3C). As anticipated, the sgRNAs for miR-16-1^mut^ and miR-15a^mut^ provoked a massive shift to the upper right and upper left quadrants, respectively (Fig. 3C). However, a small fraction of cells in which miR-15a^mut^ was targeted was found in the upper right GFP^+^dsRed^+^ quadrant. Likewise, targeting of miR-16-1^mut^ generated a significant population in the lower right GFP^+^dsRed^-^ quadrant, suggesting loss of miR-16-1^mut^ activity while retaining miR-15a^mut^ function and thus putatively refuting cluster assistance (Fig. 3C). To understand the molecular basis of these unexpected fluorescent shifts, both populations were FACS-sorted and the genomic regions corresponding to the miR-15a^mut^-miR-16-1^mut^ transgene were sequenced. This revealed that the GFP^+^dsRed^+^ population upon targeting of miR-15a^mut^ was the result of extended CRISPR/Cas9-mediated deletions, i.e. in a small subset of cells the indel mutation was not restricted to the miR-15a^mut^ locus, but included the neighboring miR-16-1^mut^ gene (Fig. S3B). Sequencing of miR-16-1^mut^ targeted GFP^+^dsRed^-^ cells revealed insertions or deletions of ≤ 2 nt within the region coding for the mature miRNA, respectively (Fig. S3C). Most likely, such indels are sufficient to disrupt the interaction of the mature miRNA with the reporter, resulting in GFP re-expression. However, we assume that the stem-loop structures carrying such restricted indel mutations still serve as substrates for the Microprocessor and thereby are capable to provide cluster assistance. Hence, these results support the finding that cluster assistance requires a processable miRNA stem-loop, and in general suggest that our CRISPR/Cas9 approach is likely to identify specific regulators of miR-15a biogenesis.

### Efficient processing of pri-miR-15a depends on SAFB2

Subsequent to the validation of the platform, we transduced screen cells with a genome-wide sgRNA library and sorted for the GFP^-^dsRed^+^ and GFP^+^dsRed^+^ populations as well as for GFP^-^dsRed^-^ cells as a reference after 8 days (Fig. S3D) (Sanjana et al., 2014). As anticipated, the analysis of sgRNAs enriched in the GFP^+^dsRed^+^ quadrant as compared to the reference revealed genes involved in general miRNA biogenesis, such as Ago2, Dgcr8, Dicer1 and Drosha as well as two positive controls targeting miR-16-1^mut^ that were spiked into the library (Table S1). Likewise, two sgRNAs targeting miR-15a^mut^ were most enriched in the GFP^-^ dsRed^+^ population, highlighting the validity of the screen (Fig. 3D and Table S2). Interestingly, within the list of > 8-fold enriched sgRNAs in this population we furthermore found all 6 sgRNAs targeting the scaffold attachment factor B2 (Safb2) gene (Fig. 3D and Table S2). SAFB2, together with SAFB1 and SLTM (SAFB-like transcriptional modulator), is part of an evolutionary highly conserved family of proteins that was originally described to bind scaffold attachment region DNA elements (Renz and Fackelmayer, 1996). However, recent studies have broadened its scope towards diverse molecular functions such as chromatin and transcriptional regulation (Alfonso-Parra and Maggert, 2010; Hernández-Hernández et al., 2013; H.-W. Liu et al., 2015; Oesterreich et al., 1997; Omura et al., 2009; Peidis et al., 2011; Yamaguchi and Takanashi, 2016), mRNA processing (Park et al., 2004; Rivers et al., 2015; Sergeant et al., 2007), the DNA damage (Altmeyer et al., 2013) and the cellular stress response (Chiodi et al., 2000; Denegri et al., 2001).

To validate a possible role of SAFB2 in miR-15a^mut^ biogenesis, we cloned and expressed individual sgRNA constructs targeting the Safb2 gene in screen cells. While control sgRNAs had no effect on reporter expression, loss of SAFB2 promoted a significant shift of cells into the dsRed^+^ only quadrant, indicating impaired miR-15a^mut^ biosynthesis (Fig. 3E). SgRNAs targeting Safb1 were not found enriched in our screen, but given that SAFB1 and SAFB2 share the same overall domain structure and are highly homologous, we also tested a potential involvement of SAFB1 in these processes. Confirming the screen, sgRNAs against Safb1 did not affect reporter expression despite being able to induce indel mutations in a T7 assay (Fig. 3E and data not shown). However, combined disruption of Safb1 and Safb2 resulted in an even stronger phenotype than of disruption of Safb2 alone, suggesting that SAFB2 is the key driver of miR-15a^mut^ biogenesis, but that SAFB1 can compensate for its loss at least to some extent (Fig. 3E). Supporting this, re-expression of human SAFB2 in sorted dsRed^+^ Safb1/2 double-knockout (DKO) screen cells was able to completely rescue the phenotype based on reporter fluorescence, whereas SAFB1 re-expression only partially restored reporter repression (Figs. 3F and G). We therefore decided to focus on SAFB2 for the remainder of this manuscript.

To ensure that SAFB2 is not only involved in the processing of the artificial miR-15a^mut^-miR-16-1^mut^ cluster overexpressed in our screen, but indeed regulates endogenous miR-15a biogenesis, we generated Safb1/2 DKO Ramos cells and analyzed them for mature miR-15a and miR-16 levels by qPCR. In accordance with the screen analysis, combined depletion of SAFB1 and 2 resulted in a significant decrease in mature miR-15a, whereas reconstitution of a CRISPR-resistant SAFB2 was able to rescue miR-15a expression to control levels (Fig. 3H). We furthermore quantified the primary miR-15a processing intermediate generated upon pri-miR-16-1 cleavage and observed its accumulation in Safb1/2 DKO cells (Figs. S3E and 3I). Again, reconstitution of SAFB2 rescued the phenotype, implying that it is critical for proper pri-miR-15a to pre-miR-15a cleavage once pri-miR-16-1 has been processed (Fig. 3I).

### SAFB2-mediated assistance is required for the productive processing of several clustered miRNAs

While our experiments clearly identify SAFB2 as a necessary component for efficient cleavage of pri-miR-15a, it is not possible to discriminate whether SAFB2 is indeed required for cluster assistance, i.e. linking miR-15a to miR-16-1, or whether it is only involved in primary miR-15a cleavage after cluster assistance has been established by other factors. We were therefore wondering whether other clustered miRNAs might also require assistance for their efficient biogenesis, and if so, whether this processing might also depend on SAFB2. To this end, we first analyzed miR-15b-16-2, a paralog of miR-15a-16-1, hypothesizing that both clusters might be processed in a similar manner. Indeed, expression of miR-15b-16-2 resulted in a strong repression of miRNA reporters for miR-15b and miR-16, respectively, whereas disruption of the miR-16-2 stem-loop by four destabilizing mutations resulted in a significant decrease in miR-15b activity (Fig. 4A). Unlike miR-15a, however, miR-15b still retained some residual activity independent of miR-16-2, suggesting that the miR-15b-16-2 cluster only partially depends on cluster assistance (Fig. 4A). Supporting this finding, CRISPR/Cas9- mediated disruption of endogenous miR-16-2 in Ramos clones lacking miR-16-1 reduced miR-15b levels to a lesser extent than miR-16-2, implying that some residual miR-15b maturation takes place (Fig. 4B). Interestingly, miR-15b lacks the prominent unpaired region that defines the lower stem of miR-15a, but exchanging its lower stem with the one of miR-26a-2 still rescued full miR-15b activity when expressed in the absence of miR-16-2 (Figs. S4A and 4C). Moreover, swapping the loop also significantly enhanced miR-15b activity, albeit less pronounced than for the lower stem (Fig. 4C). This implies that miR-15b, in analogy to miR-15a, is a suboptimal Microprocessor substrate due to features in its lower stem and loop regions. In consequence, miR-15b relies on cluster assistance to enable its full processing. To test whether cluster assistance for miR-15b depends on SAFB2, we measured mature miR-15b levels and the pri-miR-15b intermediate in wt and Safb1/2 DKO Ramos cells. Loss of SAFB proteins induced the accumulation of the processing intermediate generated after miR-16-2 processing, and consequently resulted in a significant reduction in mature miR-15b levels (Fig. 4D and E). Again, reconstitution of CRISPR-resistant SAFB2 rescued the phenotype, i.e. processing of the intermediate was restored and mature miRNA levels increased to control level (Fig. 4D and E).

**Figure 4:**
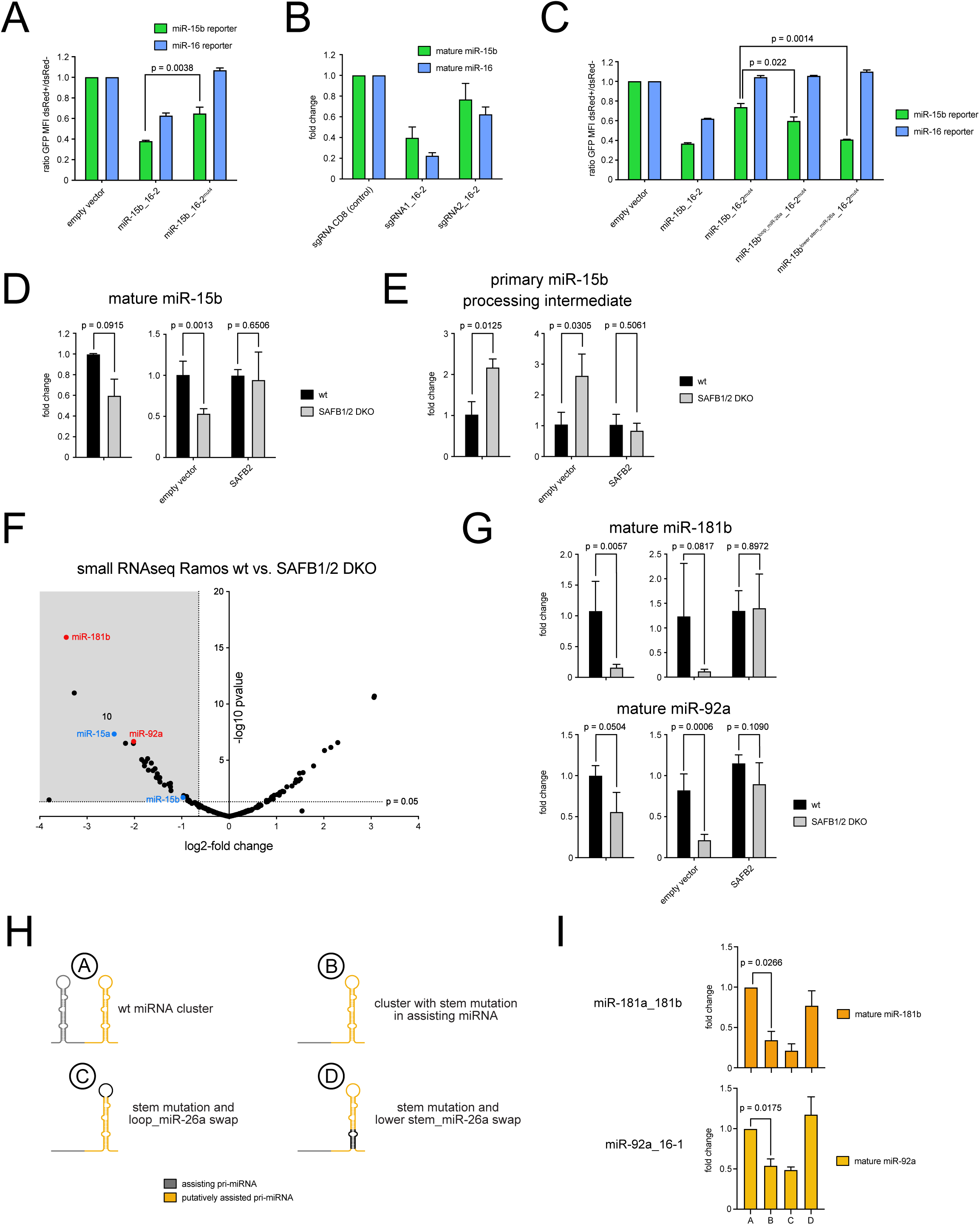
SAFB2 enables the efficient processing of several clustered miRNAs. **A**. Disruption of miR-16-2 decreases activity of the clustered miR-15b. Ramos cells expressing miR-15b and miR-16 reporters were virally transduced with constructs encoding wt miR-15b_16-2 or miR-16-2-mutated clusters and analyzed for GFP expression. **B**. Endogenous miR-15b partially depends on the neighboring miR-16-2 for its efficient biogenesis. MiR-16-1-deficient Ramos cells were lentivirally transduced with vectors encoding a CD8 sgRNA or two individual sgRNAs targeting miR-16-2. After one week, total RNA was isolated, reverse transcribed and subjected to qPCR for mature miR-15b and 16-1 levels, respectively. **C**. Exchanging the lower stem with the respective elements of miR-26a-2 abrogates the dependence of miR-15b on cluster assistance. MiRNA reporter assay using chimeric miR-15b/miR-26a_16-1 contructs that swap either the loop or the lower stem of miR-15b. **D**, **E**. SAFB2 mediates the efficient pri- to pre-miRNA cleavage of endogenous miR-15b. Quantitative RT-PCR analysis of Safb1/2 DKO and wt control cells for expression of mature miR-15b (D) or a specific processing intermediate generated upon pri-miR-16-2 cleavage (E, left panels). To rescue the phenotype, control cells and Safb1/2 DKO cells were reconstituted with an empty vector or with a vector encoding a CRISPR-resistant variant of human SAFB2. After one week, miR-15b and its processing intermediate were quantified by RT-PCR (D, E; right panels). **F**. Small RNA sequencing identifies several miRNAs that are reduced upon Safb1/2 deletion. Total RNA derived from Safb1/2 DKO Ramos cells and the corresponding controls was analyzed for global miRNA expression by small RNA sequencing. The grey square marks miRNAs that are significantly reduced in the absence of SAFB1 and 2 (reduction more than 0,7-fold (log2); p < 0.05). MiR-15a and b are shown as blue dots, miR-181b and miR-92a chosen for further analysis are depicted in red. **G**. Quantitative RT-PCR validates the SAFB2-mediated processing of miR-181b and miR-92a. Analysis of Safb1/2 DKO and control cells for mature miR-181b and miR-92a expression, respectively, under steady state conditions or upon reconstitution with SAFB2 as in D. **H**. Schematic illustration of the constructs used in Fig. 5I. **I**. MiR-181b and miR-92a rely on cluster assistance, but become independent of this effect when combined with an optimal lower stem. Quantitative RT-PCR of 293T cells expressing the indicated constructs of the miR-181a_181b and the chimeric miR-92a_16-1 cluster (Fig. S4C). Cells were transfected and analyzed in bulk after 48 hrs. MiR-181b and miR-92a levels were normalized against a spiked-in miRNA control not endogenously expressed (miR-125b). Data are represented as mean ± SD.

To investigate whether SAFB1 and 2 proteins also affect clustered miRNA expression beyond the mir-15 family, we performed small RNA sequencing in control and Safb1/2 DKO Ramos and 293T cells (Fig. 4F and S4B, Table S3). Loss of SAFB proteins resulted in reduced expression of 34 and 20 miRNAs in Ramos and 293T cells (cutoff +200 CPM; > 0,7-fold reduction; p-value < 0.05), respectively, of which 8 were downregulated in both (Table S3). Of note, all of these shared miRNAs were organized as miRNA clusters, possibly indicating SAFB-mediated cluster assistance. MiR-181b, expressed as part of the miR-181a-181b cluster, and miR-92a (miR-17-92 cluster) were chosen for further analysis. First, we confirmed their reduced expression in the absence of SAFB1 and 2 and demonstrated that their levels were restored upon reconstitution with SAFB2 (Fig. 4G). To test whether miR-181b and miR-92a were processed at least in part by cluster assistance, we then cloned the respective clusters and expressed them either in their wt form or as a variant in which the supposedly assisting miRNA was disrupted by extensive stem mutation (Fig. 4H). Since the miR-17-92 cluster encodes miR-92a and five additional miRNAs that could in principle mediate cluster assistance, we simplified the cluster organization and expressed miR-92a together with miR-16-1 or its mutated form, thereby mimicking a bicistronic cluster (Fig. S4C). Remarkably, disruption of the assisting miRNA significantly decreased mature miR-181b and miR-92a levels compared to the control setting (Fig. 4I), clearly demonstrating an involvement of cluster assistance in these SAFB-regulated miRNA biogenesis processes. Having demonstrated that both miR-15a and b are suboptimal Microprocessor substrates that require the clustered miRNA for their efficient processing, we hypothesized that this feature might be a common denominator of cluster assistance. Indeed, swapping the lower stem of the prototypic miR-26a-2 into the respective constructs rendered miR-181b and miR-92a processing independent of an additional miRNA on the same primary transcript, confirming that cluster assistance is only required for suboptimal DGCR8/DROSHA substrates (Fig. 4I). Together, these experiments demonstrate that several miRNAs depend on cluster assistance for their efficient biogenesis, and that the respective stem-loop structures serve as imperfect Microprocessor substrates as a common feature. Moreover, efficient processing of these miRNAs depends on SAFB2, strongly indicating its direct involvement in cluster assistance. This, however, raises the question how SAFB2 can license the cleavage of a suboptimal Microprocessor substrate in the presence of an additional miRNA on the primary transcript, i.e. how cluster assistance is mediated mechanistically.

### SAFB2 binds to the microprocessor

In accordance with their proposed role as regulators of gene expression and RNA processing, SAFB proteins contain a conserved RNA recognition motif (RRM, Fig. 5A) and have been identified as RNA binding proteins by two unbiased studies (Baltz et al., 2012; Castello et al., 2012). We therefore hypothesized that SAFB2 might employ its RNA binding capacity to directly associate with the primary miRNA structures that depend on cluster assistance, thereby enabling efficient Microprocessor cleavage. However, mutants of SAFB2 lacking the RMM or the arginine/glycine motif RGG/RG, which has been shown to be sufficient for RNA binding in SAFA (scaffold attachment factor A; (Kiledjian and Dreyfuss, 1992)), were as potent as full-length SAFB2 in restoring miR-15a^mut^ reporter repression in Safb1/2 DKO screen cells (Fig. 5B). This implied that other domains of SAFB2 are required to mediate cluster assistance, and motivated us to test a larger panel of SAFB2 mutants in this experimental setup (Fig. 5C). Surprisingly, most domains of SAFB2 turned out to be dispensable for reporter repression. In fact, only the deletion of the regions between the RRM and the C-terminal RG-rich domain significantly impaired or abolished cluster assistance (Fig. 5C, upper panel). To investigate whether this part of SAFB2 was not only necessary, but possibly also sufficient, we furthermore tested incrementally smaller versions of SAFB2 for their potential to rescue the knockout phenotype. Here, we found that a SAFB2 fragment comprising amino acids 561 to 726 (hereafter termed SAFB2^min^), which encodes a putative coiled-coil domain and the arginine/glutamic acid-rich (RE) region, was almost equally active as the full-length protein (Fig. 5C, lower panel). Interestingly, the function of SAFB2^min^ appeared to be highly conserved within chordates, as expression of the corresponding SAFB^min^ region of the lancelet *Branchiostoma floridae* was also able to rescue reporter repression, whereas *Drosophila melanogaster* SAFB^min^ was not (Fig. S5A and S5B).

**Figure 5:**
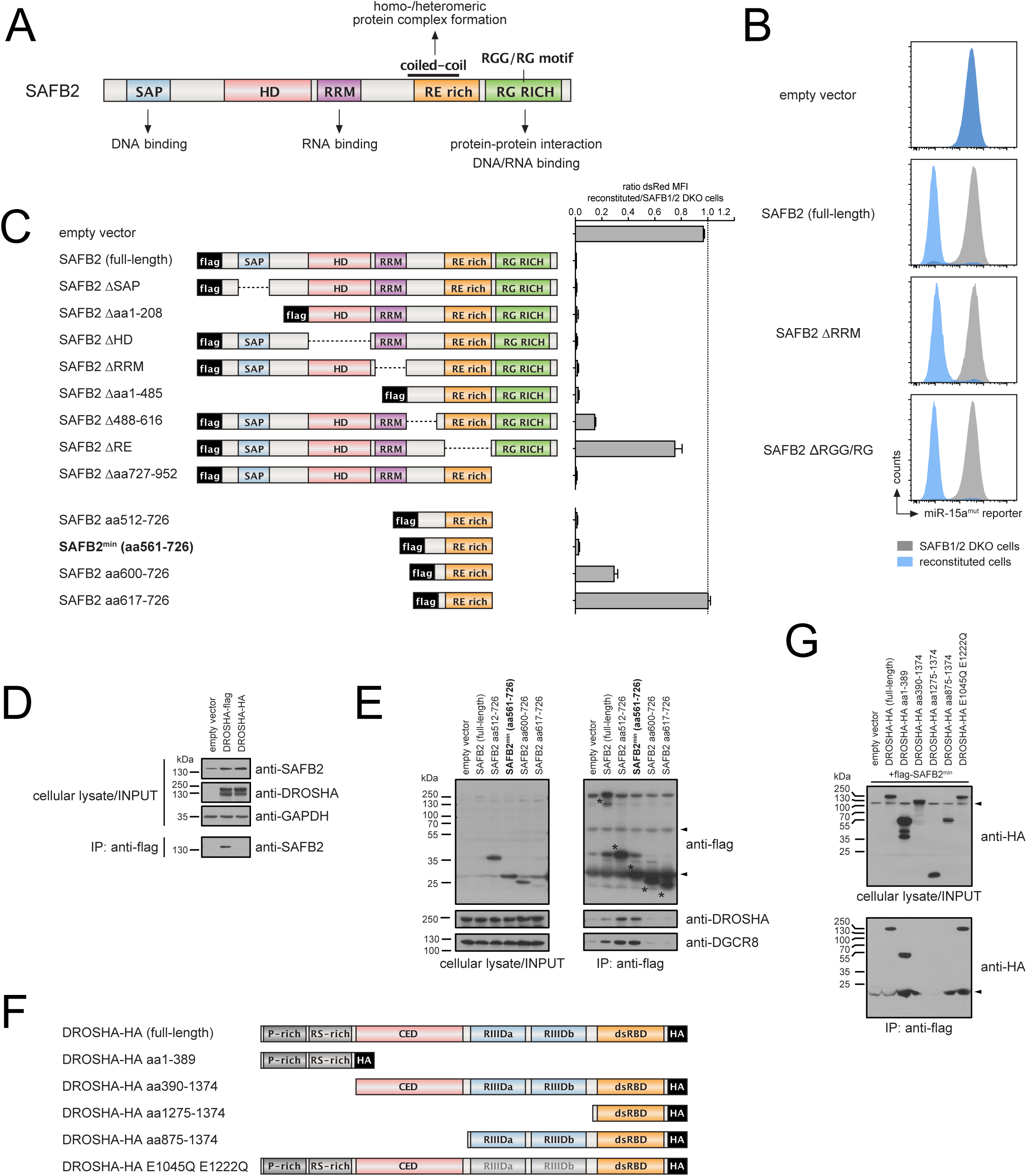
SAFB2 associates with the Microprocessor. **A**. Schematic illustration of the domain structure and their putative functions for human SAFB2 (not in scale). SAP, *SAFA/B, Acinus and PIAS*; HD, *homologous domain* in SAFB1 and 2; RRM, *RNA recognition motif*. **B**. SAFB2 function in miR-15a^mut^ processing is independent of its canonical RNA binding. Screen cells lacking SAFB1 and 2 and sorted for the dsRed+ population were reconstituted with full-length SAFB2 or with mutants lacking the RNA recognition (ΔRRM) or the RGG/RG motif. Non-transduced cells are shown as a grey histogram, transduced cells are depicted in blue. **C**. The described SAFB2 function in miR-15amut processing can be linked to a small part of comprising the putative coiled-coil domain. Screen cells as in B were transduced with the depicted deletion mutants and analyzed for miR-15a^mut^ reporter expression by flow cytometry after one week. The individual bars show the ratio of dsRed MFI comparing transduced versus non-transduced cells. A ratio of 1 representing no effect is marked by a dashed line. **D**. SAFB2 associates with the Microprocessor. DROSHA-deficient 293T cells were transfected with a control vector or with vectors encoding either flag-tagged or HA-tagged DROSHA. After 48 hrs, cells were lysed and subjected to anti-flag immunoprecipitation. Western blot analysis shows DROSHA and GAPDH expression in the input as well as SAFB2 in input and the IP. **E**. SAFB2^min^ precipitates endogenous DROSHA. Inversely to D, 293T cells transfected with constructs encoding full-length flag-SAFB2, flag-SAFB2^aa512-726^, flag-SAFB2^min^ or further shortened SAFB2 mutants were lysed and analyzed for their association with the Microprocessor. Probing the input and the IP with anti-flag antibodies served as a control. SAFB2 fragments are marked by black asterisk. Light and heavy chain of the precipitating antibody in the IP are marked by black triangles. **F**. Schematic illustration of DROSHA deletion mutants. **G**. SAFB2 binds to the N-terminus of DROSHA. DROSHA-deficient 293T cells were co-transfected with a vector encoding flag-SAFB2^min^ and different DROSHA mutants as shown in F. After 48 hrs, cellular lysates were subjected to anti-flag immunoprecipitation and analyzed for co-purification of HA-tagged DROSHA. Unspecific bands are marked by black triangles.

Due to the lack of any canonical RNA binding domain in SAFB2^min^, we hypothesized that SAFB2’s activity in miRNA processing is mediated by protein-protein interaction. In particular, we were wondering whether SAFB2 is directly binding to the Microprocessor, given that this enzymatic complex acquires the ability to cleave suboptimal substrates in the context of cluster assistance. Indeed, immunoprecipitation of tagged DROSHA in Ramos cells revealed its association with SAFB2, and vice versa, immunoprecipitation of full-length SAFB2, SAFB2^aa512-726^ or the even shorter SAFB2^min^ pulled-down DROSHA and DGCR8 (Figs. 5D and E). Further shortening the N-terminal part of SAFB2^min^ abolished this interaction (Fig. 5E), and such shortened SAFB2 mutants failed to restore full miR-15a reporter repression in Safb1/2 DKO screen cells (Fig. 5C, lower panel). This strongly implies that the interaction between SAFB2 and the Microprocessor is a prerequisite for cluster assistance. To map this interaction in more detail, we performed pull-down experiments with different DROSHA fragments and observed strong binding of SAFB2^min^ to the N-terminal part of DROSHA (aa1-389), but not to the structurally resolved C-terminal part (aa390-1374) (Fig. 5F). This suggests that SAFB2 associates with DROSHA, either directly or indirectly, and not with DGCR8, since the DGCR8-DROSHA interface has been mapped to the DROSHA RNAse III domains (aa876-1056 and aa1107-1233). Together, these data indicate that SAFB2 exerts its function in cluster assistance in an RNA binding-independent manner, but rather through association with the Microprocessor.

## Discussion

Many of the sequence and structural features that define a primary miRNA stem-loop and thereby allow the discrimination of authentic miRNAs from similar short RNA stem-loop structures have been identified in recent years (Auyeung et al., 2013; Fang and Bartel, 2015; Han et al., 2006; Lee et al., 2003; Zeng et al., 2005; Zeng and Cullen, 2005). However, we are still far from a complete understanding of rules that govern cleavage of primary miRNAs, a process that ultimately defines the mature miRNA pool. This especially applies to the recently described phenomenon of co-dependent miRNA expression within clusters, in which the Microprocessor-mediated processing of certain miRNAs appears to be linked to the presence of a second miRNA stem-loop on the same primary transcript (Haar et al., 2016; Lataniotis et al., 2017; Truscott et al., 2016).

Here, we have added the well-known tumor suppressor miR-15a to the growing list of miRNAs that require cluster assistance for their efficient processing and used extensive mutagenesis of the miR-15a-16-1 cluster to define the molecular features underlying this phenomenon. Our data demonstrate that pri-miR-15a is a poor Microprocessor substrate when expressed on its own, as it failed to efficiently recruit DROSHA due to a large unpaired region within its basal stem. Previous studies have already shown the importance of extensive pairing at the base of the miRNA stem-loop (Han et al., 2006; Zeng and Cullen, 2003), and this has been confirmed by recent high throughput analyses (Fang and Bartel, 2015). Accordingly, mutations that increase the extent of base pairing within the miR-15a lower stem reduced or even abrogated its dependence on miR-16-1. Although only indirectly, our data indicate similar imperfections for clustered miR-15b, miR-181b and miR-92a, all of which showed strongly reduced expression in the absence of an assisting miRNA. However, processing of these primary miRNAs was rescued by swapping their lower stem regions to the one of prototypic miR-26a, suggesting that sequence and/or structural features within this region are responsible for their subpar cleavage. It is therefore tempting to speculate that all miRNAs that rely on cluster assistance for their efficient processing are suboptimal Microprocessor substrates. Supporting this hypothesis, miR-998 and miR-125, both of which had been previously identified to depend on cluster assistance in *Drosophila*, are characterized by either an unusually large loop of >60 nt or an extended lower stem, respectively (Truscott et al., 2016). Along the same line, structural analyses of the Epstein-Barr virus primary miR-BHFR1-3, which is barely expressed without the assistance of its neighboring miR-BHFR1-2, revealed that only a small fraction of transcripts formed a miRNA stem-loop when folded *in vitro* (Haar et al., 2016).

This raises the question how such suboptimal substrates can be processed in the first place. Our data confirm that cluster assistance does not require the specific function of the clustered miRNA, but is rather mediated by any additional stem-loop that can be cleaved by the Microprocessor as a *cis* regulatory element (Haar et al., 2016; Truscott et al., 2016). In line with previous data, we demonstrate that both the relative position and the distance of the clustered miRNA stem-loops appear irrelevant within the tested parameters (Truscott et al., 2016), suggesting that the Microprocessor may only require to be recruited to any position of the primary transcript in order to orchestrate the co-dependent cleavage of stem-loop structures. However, we speculated that additional factors might be involved in cluster assistance, and indeed identified the SAFB family, in particular SAFB2, as an important element for pri-miR-15a processing. Interestingly, cluster assistance appears to be independent of canonical DNA and RNA binding, but rather appears to be mediated by either direct or indirect interaction of SAFB2 with DROSHA. Such an RNA-binding independent function of SAFB2 seems to contradict recent CLIP data that suggest the direct association of SAFB1 with a number of primary miRNAs, among them the transcript that comprises the miR-17-92 cluster (Rivers et al., 2015). In this study, concomitant knockdown of SAFB1 and SAFB2 in neuroblastoma cells led to a decrease in mature miR-19a, but not miR-17 or 18 encoded by the same cluster. Notably, these changes were accompanied by an increased expression of the primary transcript, pointing to a role of SAFB proteins in pri-miRNA processing that actually does involve direct RNA binding (Rivers et al., 2015). In our system, we observed that both SAFB1 and SAFB2 contribute to miR-15a^mut^ processing, but we cannot exclude that they function by different mechanisms. It might very well be that SAFB1 binds to pri-miRNA, whereas SAFB2 associates with the Microprocessor and does not rely on its RNA-binding property to enable cluster assistance. In fact, it will be interesting to decipher how SAFB1 and SAFB2 differ that much in their ability to mediate miRNA processing, given their high sequence similarity (Oesterreich et al., 1997; Renz and Fackelmayer, 1996; Townson et al., 2003). Of course, deletion of either SAFB1 or SAFB2 in mice already revealed non-redundant phenotypes, with high lethality, severe growth retardation and male infertility in the former, but no major defects in the latter (Ivanova et al., 2005; Jiang et al., 2015). Thus, both family members indeed appear to exert different molecular functions.

With respect to SAFB2, we were able to pinpoint its activity in cluster assistance to a relatively small but highly conserved part of the protein that comprises the arginine/glutamic acid-rich (RE) region and a short stretch upstream of this domain. How do SAFB2 and SAFB2^min^ enable pri-miR-15a, -15b, -181b and 92a cleavage in a clustered setup? Our data indicate that loss of the SAFB2:DROSHA complex correlates with a loss of function in the fluorescent reporter system. Hence, we postulate that binding of SAFB2 to DROSHA is indeed a key step in cluster assistance. Notably, the minimally functional region of the SAFB2 protein in terms of miRNA processing also contains a putative coiled-coil domain, a structure that has been associated with various functions, among them the formation of homo- and heteromeric protein complexes (Wang et al., 2012). It is therefore tempting to speculate that SAFB2 complexes promote the dimerization of the Microprocessor, which might allow the processing of suboptimal substrates by one enzymatic unit once the other DROSHA/DGCR8 complex has been recruited to the assisting miRNA stem-loop. Alternatively, binding of SAFB2 may induce a conformational shift or stabilize a certain conformation in the central domain of DROSHA, which is the region that binds to the lower stem of pri-miRNAs miRNAs (Kwon et al., 2016; Nguyen et al., 2015). These structural changes may be sufficient to alter the specificity of the Microprocessor complex towards imperfect substrates once it has been recruited to the primary miRNA through a near-perfect stem-loop structure. Third, it is conceivable that the Microprocessor remains bound to the primary transcript after the assisting stem-loop has been cleaved, and that it then moves alongside the RNA to scan for additional substrates in a SAFB2-dependent manner. Again, this would overcome the necessity for direct DROSHA/DGCR8 recruitment, enabling the processing of alternative stem-loops.

With respect to its biological role, it is conceivable that cluster assistance serves a regulatory function, possibly coordinating the spatial and/or temporal expression of miRNAs with complementary function. However, we favor a model in which cluster assistance simply expands the cellular miRNA repertoire by rendering suboptimal stem-loop structures accessible for Microprocessor-mediated cleavage, thereby allowing the formation of additional or more complex gene regulatory networks. Notably, both the minimally functional domain of SAFB2 as well as structural RNA features, i.e. the unpaired region within the lower stem of pri-miR-15a, are highly conserved, indicating that cluster assistance has been a beneficial evolutionary acquisition.

In conclusion, our study reveals a critical function of cluster assistance in primary miRNA processing and indicates that a neighboring miRNA stem-loop can compensate in *cis* for structural defects in suboptimal Microprocessor substrates. Thus, cluster assistance appears to have the same effect as structural or sequence-specific features within the stem-loop itself, such as the UG, the CNNC or the mismatched GHG motifs (Fang and Bartel, 2015). Mechanistically, our findings suggest a model in which SAFB2 enables the cleavage of imperfect DGCR/DROSHA substrates in a clustered setup, but it needs to be determined how exactly SAFB2 exerts its function. With the limited number of pri-miRNA stem-loops that have been identified to rely on cluster assistance, it appears difficult to clearly define specific features for this type of regulation and, based on that, to extrapolate which of the large number of clustered miRNAs in the human genome will be regulated likewise. In the future, it will therefore be crucial to identify those miRNAs and their common denominatory features. Given the broad structural and sequence varieties of clustered miRNAs, it is unlikely that SAFB proteins are the only factors assisting suboptimal substrate processing. In fact, concomitant loss of SAFB1 and SAFB2 in our experiments invoked a much lesser effect on pri-miR-15a processing than disruption of miR-16-1, suggesting that already for this cluster other, thus-far unknown factors are involved as well. It is tempting to speculate that additional DROSHA and/or DGCR8-accessory factors may enable cluster assistance, performing the same function yet for different miRNA families.

## Methods

### Cells, cell culture and manipulation

The Burkitt lymphoma line Ramos (DSMZ #ACC603) and the murine B cell progenitor Baf3 (DSMZ #ACC300) were cultured in RPMI medium (Sigma) supplemented with 7.5% fetal calf serum, L-glutamin, 100 U/ml penicillin, 100 U/ml streptomycin (all Sigma) and WEHI-3B cell derived IL-3 in excess (Baf3 only; Ref. PMID: 6122701). 293T cells (DSMZ #ACC635) were cultured in IMDM medium with the same supplements. For the generation of retro- or lentiviruses, 293T cells were transfected with DNA plasmids of interest and with either pSPAX2 or HIT60 together with VSVg at a 4:1:1 ratio. DNA was introduced into the cells using PEI (polyethylenimine, Polysciences) at a 3:1 PEI:DNA ratio. Viral supernatants were harvested after 36 and 60 h, mixed with polybrene (8 to 16 µg/ml final concentration) and used for spin-infection of cells in 1,5 ml reaction tubes or 96-well plates (400 g at 37°C for 60-90 min).

### MiRNA activity assay and Microprocessor assay

To quantify miRNA activities, Ramos or 293T cells expressing GFP-based fluorescent miRNA reporters were retrovirally transduced with miRNA vectors. Reporter repression was quantified by calculating the ratio of GFP mean fluorescence intensity (MFI) in transduced dsRed^+^ cells compared to the non-transduced dsRed^-^ population. If not stated otherwise, individual values were normalized based on cells expressing an empty control vector.

To measure the activity of the Microprocessor *in vivo*, we adopted and modified a previously published protocol (Allegra and Mertens, 2011). In short, the stem-loop structures or mutants thereof were cloned into the region corresponding to the 3’-UTR of a GFP cDNA. Transduction of these contructs at a low multiplicity of infection (MOI) gave rise to GFP-positive cells of a defined MFI which is inversely correlated with Microprocessor activity.

### Proximity biotinylation/RaPID

RNA-protein interaction detection (RaPID) in living cells was performed according to a recent publication (Ramanathan et al., 2018). In short, 293T cells were cotransfected with vectors encoding an RNA bait flanked by BoxB stem loops and the BASU biotin ligase (Addgene #107253 and #107250, kindly provided by Paul Khavari) at a 4:1 ratio. After 48 hrs in normal growth medium, cells were further cultered with 200 µM biotin (final conc.; Sigma) for 4 hrs before lysis in RIPA buffer (50 mM Tris HCl, pH 7.4, 1% NP-40, 0.25% sodium deoxycholate, 150 mM NaCl, 1 mM EDTA (pH 8), protease inhibitor cocktail; Sigma). Streptavidin-agarose beads (GE healthcare) were then incubated with the cleared lysates over night at 4°C, followed by consecutive washing steps according to the protocol (Ramanathan et al., 2018). Finally, beads were boilded in reducing sample buffer containing 4 mM biotin for 10 min and the supernatant was subjected to SDS-PAGE and western blotting onto PVDF membranes.

### Plasmids

For the expression of human primary miRNAs, the respective cassettes were cloned from genomic DNA and ligated into a modified LMP-dsRedExpress2 (reference) in exchange for the miR-30-based shRNA cassette. In case of miR-15a^mut^_16-1^mut^ used in the CRISPR screen, the cluster was cloned 3’ of a cDNA encoding a tailless CD8 as a marker into an MSCV-based retroviral backbone derived from pMIG (reference).

Reporter constructs were generated by oligonucleotide annealing and cloning of the resulting DNA cassettes into LMP-destGFP (Knackmuss et al., 2015), 3’ of a cDNA encoding a destabilized GFP-PEST fusion. SAFB1 and SAFB2 and their mutants were expressed as flag-tagged proteins from pMIG-derived pMITHy1.2, in which GFP was exchanged for cell surface marker Thy1.2. DROSHA expression constructs were based on pCR-DROSHA-flag (kindly provided by N. Kim; reference). Mutagenesis of all constructs was performed by overlapping extension PCR.

For CRISPR/Cas9-mediated gene deletion, cells were lentivirally transduced with CRISPR_V2-based vectors (Addgene #52961; supp. Materials; (Sanjana et al., 2014)), selected with 2 µg/ml puromycin and analyzed after 8 days or plated in methycellulose to isolate single cell clones.

### CRISPR/Cas9 screen

Screen cells were generated by transduction of parental Baf3 cells with constructs encoding the GFP-based miR-16^mut^ and the dsRed-based miR-15a^mut^ reporters, followed by the mutated miR-15a^mut^_16-1^mut^ cluster and a construct encoding Cas9 (Addgene #52962).

For virus production, 293T cells seeded onto 150mm dishes were transfected with 21 µg of the murine lenti-GeCKO-V2 plasmid library (Addgene pooled library #1000000053), 0.168 ng each of the sgRNAs targeting miR-15^mut^ and miR-16-1^mut^, 15.75 µg of pSPAX2 and 5.25 µg of VSVg plasmids. Viral supernatant was harvested after 36h and 50h, concentrated by Amicon ultra-15 centrifugation tube (Merck), snap frozen in liquid nitrogen and stored at −80°C.

The CRISRP/Cas9 screen was performed according to a protocol adapted from a previous study (Shalem et al., 2014). In short, for each replicate 1 x 10^9^ Baf3 screen cells were transduced at an MOI of about 0.25 and selected with puromycin (2µg/ml) after 24h. After 8 days in culture, screen cells were harvested and FACS sorted for dsRed^+^GFP^-^, dsRed^+^GFP^+^ and about 2,5 x 10^7^ dsRed^-^GFP^-^ cells as a reference using an ARIA III (Becton Dickinson). Genomic DNA was isolated using a commercial kit (Macherey-Nagel) and sgRNA sequences were amplified by a nested PCR approach (see suppl. materials section for DNA oligo sequences) that added stagger bases, specific barcodes and the illumina adapters. The PCR products were separated on a gel, purified by agarose gel extraction, quantified and then pooled before sequencing on a Hiseq4000 (Illumina) in collaboration with the Biomedical Sequencing Facility (BSF, Vienna, Austria). Sequencing data were analyzed with the publically available online tools GenePattern (Reich et al., 2006) and Galaxy (Afgan et al., 2018). Reads were first demultiplexed and trimmed followed by alignment of the sgRNA sequences to a reference using Bowtie2 (Langmead and Salzberg, 2012) (see Table S4 for detailed sgRNA counts for all experimental replicates). SgRNAs enriched in the dsRed^+^GFP^-^ and dsRed^+^GFP^+^ populations were identified using the edgeR shRNAseq tool (Dai et al., 2014).

### SmallRNA sequencing

Libraries for Illumina sequencing were prepared with the Somagenics RealSeq-AC miRNA library Kit, using 100 ng total RNA isolated with the miRNEasy system (QIAGEN) as input. The samples were individually indexed and the libraries were sequenced on Illumina HiSeq 3000/4000 instruments in 50-base-pair-single-end mode at the Biomedical Sequencing Facility (BSF) of the Research Center for Molecular Medicine (CeMM), Vienna. Base calls provided by the Illumina Real-Time Analysis (RTA) software were subsequently converted into unaligned BAM format for long-term archival and de-multiplexed into sample-specific BAM files via custom programs based on Picard tools (Ref: http://broadinstitute.github.io/picard/). For data analysis, sequencing-adapters were trimmed with TrimGalore and reads were mapped against a reference of all mature human miRNAs (miRbase.org) using HISAT2 (Kim et al., 2015; Kozomara and Griffiths-Jones, 2011). Differential expression analysis of control and knockout cell lines was performed with edgeR (R. Liu et al., 2015; Robinson et al., 2010). Sequencing data have been deposited under GEO accession number GSE141098.

### Quantitative RT-PCR

For quantification of mature miRNAs, total RNA was isolated with the miRNeasy system (Qiagen) or the Universal RNA Purification kit (Roboklon). RNA samples were reverse transcribed into cDNA and analyzed by PCR using either the Taqman MiRNA Reverse Transcription kit, Taqman miRNA assays and probe-based PCR master mix (Vazyme) or the Qiagen MiRCURY LNA RT system followed by the respective miRNA assay system and SYBR PCR master mix (Vazyme) according to the manufacturer’s protocol. As a control, miRNA expression was normalized against housekeeping snoRNA-202 or miR-16. The primary miRNA processing intermediates for miR-15a and 15b as well as mature miR-181a and b were quantified by self-designed stem-loop assays according to a previous protocol (Varkonyi-Gasic et al., 2007). Oligonucleotide sequences are available in the suppl. Materials section. Fold changes were calculated with the 2^(-ΔΔC(t))^ method (Livak and Schmittgen, 2001).

### Northern blot

Northern Blot analysis was performed as previously described (Runte et al., 2001). Briefly, 10 µg of total RNA was size separated on an 18% denaturing polyacrylamide gel and subsequently blotted onto an Amersham HybondTM-N+ membrane (GE Healthcare) and cross-linked. MicroRNAs miR-15a and miR-16-1 were detected with 10 pmol of 5’-radioactively labeled, 2’O-methylated probes, respectively (suppl. Material). Signal was detected after 3 days, using a Typhoon FLA 9500 (GE Healthcare).

### Immunoprecipitation and Western blotting

About 1 x 10^6^ to 2 x 10^7^ cells were lysed in ice-cold RIPA buffer (50 mM Tris HCl, pH 7.4, 1% NP-40, 0.25% sodium deoxycholate, 150 mM NaCl, 1 mM EDTA (pH 8), protease inhibitor cocktail; Sigma). For immunoprecipitations, 1/20 of each sample was saved as an input control and the remaining lysate was incubated with anti-flag M2 agarose beads (Sigma) over night at 4°C on a turning wheel. Beads were washed 4 times with lysis buffer and together with the total cellular lysate were mixed with reducing sample buffer containing 100 mM DTT. Samples were boiled for 6 min and subjected to SDS-PAGE and western blotting onto PVDF membranes. Proteins were detected using primary antibodies against SAFB1, SAFB2, DGCR8 (all Bethyl), DROSHA, beta-actin, GAPDH (all Cell Signaling Technologies), HA and flag epitope tags (Roche and Sigma). Immunoreactive proteins were visualized with HRP-labeled secondary antibodies (Pierce) and the ECL system (Advansta) on light-sensitive film.

### Flow cytometry

Single-cell suspensions were stained using anti-CD8a (Biolegend) and anti-Thy1.2 (Biolegend) followed by streptavidin-APC (eBioscience). Data were acquired on an LSR Fortessa or an Aria Sorter (Becton Dickinson).

### Statistics

Statistical significance was calculated with paired (Figs. 1B, 1C, S1B, 2B, 2E, 4A and 4E) or unpaired (Figs. 3H, 3I, 4D and 4E) t-tests when only two groups were compared, or with one-way ANOVA followed by Tukey post-hoc tests for more than two groups (Figs. 2D and 4C). For quantitative PCRs, data are presented as fold changes compared to a control sample to simplify visual interpretation. However, statistical significance was calculated on the ΔCt values of the respective samples since fold-changes are not normally distributed.

### Software

Statistical analysis was performed using Prism 5 and 8 (Graphpad) and flow cytometric data were analyzed and visualized using Flowjo (Becton Dickinson). Figure were prepared with Affinity Designer (Serif). RNA secondary structures were predicted with the RNA-fold web server (Lorenz et al., 2011) and visualized with Varna (Darty et al., 2009). Sequence alignments were executed and visualized with Clustal Omega and ClustalW2 (Madeira et al., 2019).

## Supporting information

Differential miRNA expression in SAFB1/2 knockout vs. control cells

CRISPR/Cas9 screen: Total sgRNA counts in sorted cell populations for all experimental replicates

CRISPR/Cas9 screen: Enriched sgRNAs in GFP+dsRed+ as well as GFP-dsRed+ populations

## Acknowledgements

We thank C. Soratroi and I. Gaggl for their technical assistance, D. Bartel for discussions and the staff of the Biomedical Sequencing Facility of the Research Center for Molecular Medicine, Vienna, for advice and sequencing services. This study was supported by the Austrian Science Fund (FWF; grants P-30194 and P-30196) to S.H.. M.L. is a recipient of a DOC PhD fellowship by the Austrian Academy of Science (ÖAW).

## Author contributions

Conceptualization, S.H.; Methodology, K.H., M.L., S.A. and S.H; Formal Analysis, M.L.; Investigation, K.K, M.L., A.J., S.A., F.E., S.M.H. and S.H.; Writing – Original Draft, S.H.; Writing – Review and Editing, A.V., V.L., A.H. and S.H.; Visualization, K.H., M.L. and S.H.; Supervision, V.L., A.H. and A.V.; Funding Acquisition, A.V. and S.H.

## Supplemental Information

Table S1: CRISPR screen; enriched sgRNAs in the dsRed^+^GFP^+^ population

Table S2: CRISPR screen; enriched sgRNAs in the dsRed^+^GFP^-^ population

Table S3: smallRNA seq; analysis for differentially regulated miRNAs in control vs. Safb1/2 DKO Ramos and 293T cells

Table S4: CRISPR screen; mapped sgRNA read counts for all experimental replicates

Suppl. material: Detailed sequence information of all constructs used in this study.

---

**Figure S1.**
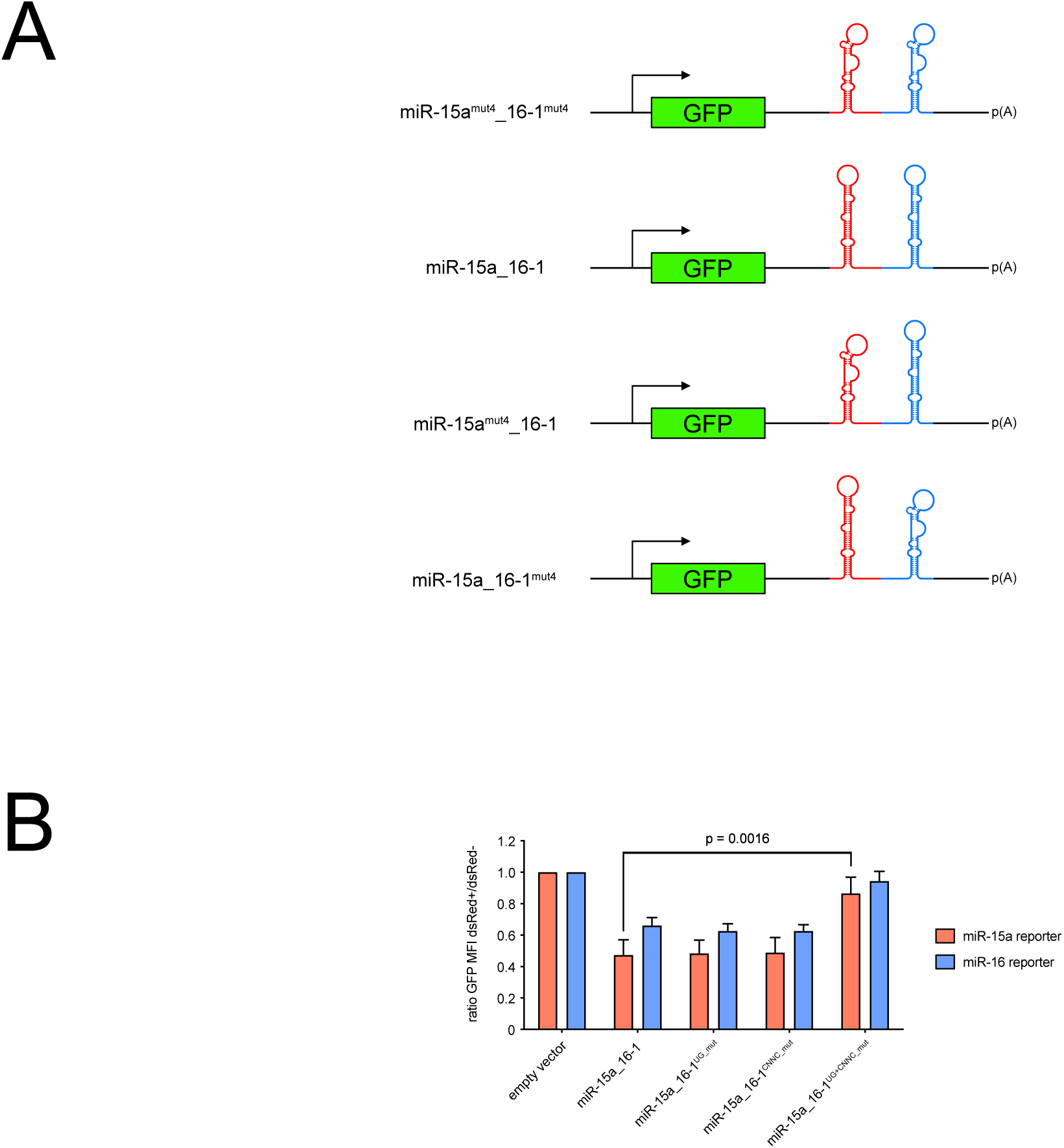
**A**. Schematic illustration of the constructs used for the Microprocessor assay. In short, cassettes encoding the miR-15a_16 cluster or variants thereof in which either one or both stem-loops have been deleted by mutagenesis are cloned into the 3’-UTR of a GFP cDNA. When expressed within cells, Microprocessor-mediated processing of the transcript cleaves off the polyA tail, resulting in reduced mRNA stability and consequently low GFP expression. In consequence, the GFP intensity is reciprocally proportional to the Microprocessor activity. **B**. Primary miR-15a cleavage depends on the processing of pri-miR-16-1. Ramos cells expressing miR-15a and miR-16 reporters were transduced with constructs encoding the wt miR-15a_16-1 cluster or UG and/or CNNC motif mutants of miR-16-1. GFP expression was analyzed after 48 hrs and plotted as the ratio of GFP MFI comparing the transduced dsRed+ cells with the non-transduced dsRed-cells. Values were normalized to the empty vector control.

**Figure S2.**
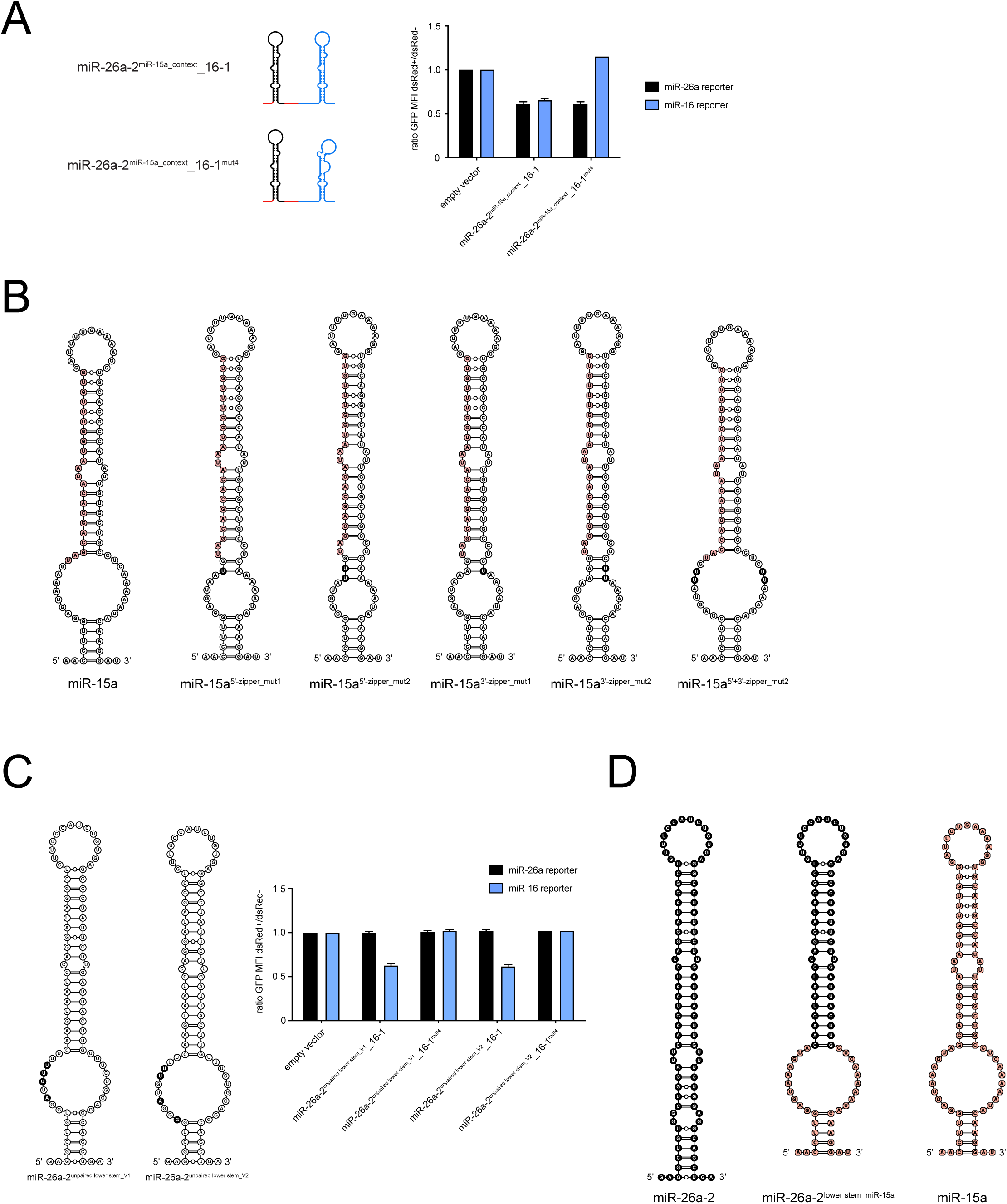
**A**. A miR-26a-2 stem-loop comprising the apical loop, upper and lower stem is processed in the miR-15a context even in the absence of miR-16-1. Ramos cells sorted for miR-26a and miR-16 reporter expression were transduced with the depicted constructs and analyzed for reporter repression by flow cytometry after 48 hrs. The bar graph shows the ratio of GFP MFI values comparing transduced and non-transduced cells, normalized to the empty vector control. **B**. Schematic illustration of the predicted structures of miR-15a constructs and mutants used in Fig. 2D. Nucleotides corresponding to the mature miRNA are drawn in red, mutated nucleotides in black. **C**. Random unpairing mutations in the lower stem disrupt miR-26a-2 function independent of cluster assistance. Ramos cells expressing miR-26a or miR-16 reporters were transduced with lower stem mutants as depicted (mutated nucleotides are drawn in black) and tested for GFP expression after 48 hrs. Individual bars show the GFP MFI ratio of dsRed+/dsRed- cells normalized to the control. D. Schematic illustration of the predicted structures of the chimeric miR-15a/26a-2 construct (center) and its two origins. MiR-26a-2-derived nucleotides are shown in black, miR-15a-derived nucleotides in red.

**Figure S3.**
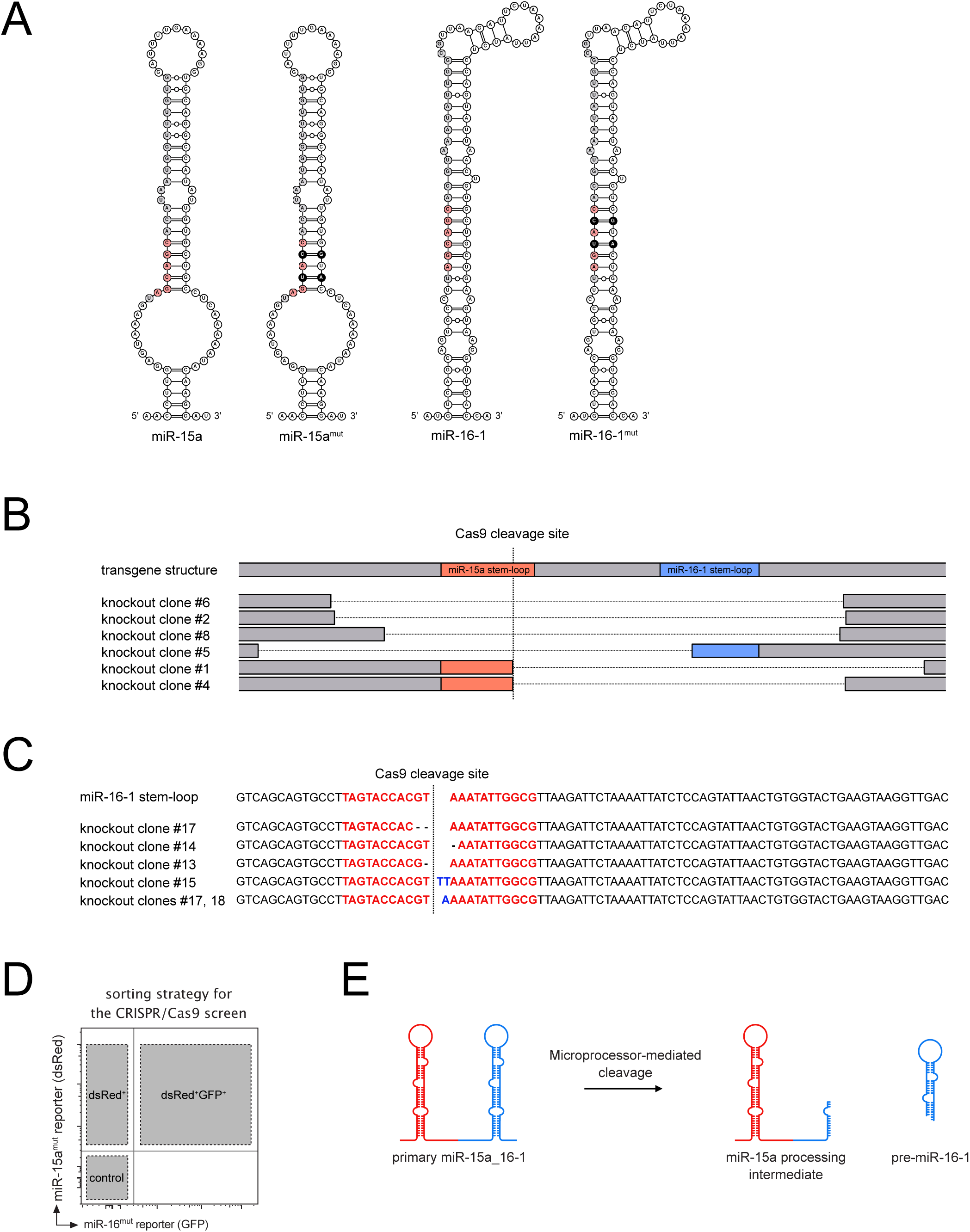
A. Schematic illustration of the predicted structures of wt miR-15a and 16-1 as well as the corresponding miR-15^mut^ and miR-16-1^mut^ variants used for the CRISPR/Cas9 screen. The seed regions are colored in red, the mutated nucleotides are marked as black. **B**. Cas9-mediated cleavage and repair of the miR-15a^mut^ gene often extends into the neighboring miR-16-1^mut^ gene. Schematic illustration of transgene DNA sequences derived from sorted dsRed+GFP+ screen cells expressing an sgRNA targeting miR-15a^mut^. A dashed line marks the predicted cleavage site. Compared to the control, all analyzed clones displayed large genomic deletions that disrupted miR-16-1mut expression. **C**. DsRed-GFP+ screen cells upon Cas9-mediated cleavage and repair of the miR-16-1^mut^ gene show a restricted pattern of indel mutations. Transgene DNA sequences of the miR-16-1^mut^ gene derived from sorted dsRed-GFP+ screen cells expressing an sgRNA targeting the locus. A dashed line marks the predicted cleavage site, the sequence corresponding to the mature miRNA is depicted in red. Compared to the control, the knockout clones show deletions (-) or insertions (blue letters) of only one to two nucleotides. **D**. Sorting scheme for the CRISPR/Cas9 screen. **E**. Illustration of the putative sequential processing of the primary miR-15a_16-1 cluster and the resulting miR-15a processing intermediate that can be quantified by quantitative PCR.

**Figure S4.**
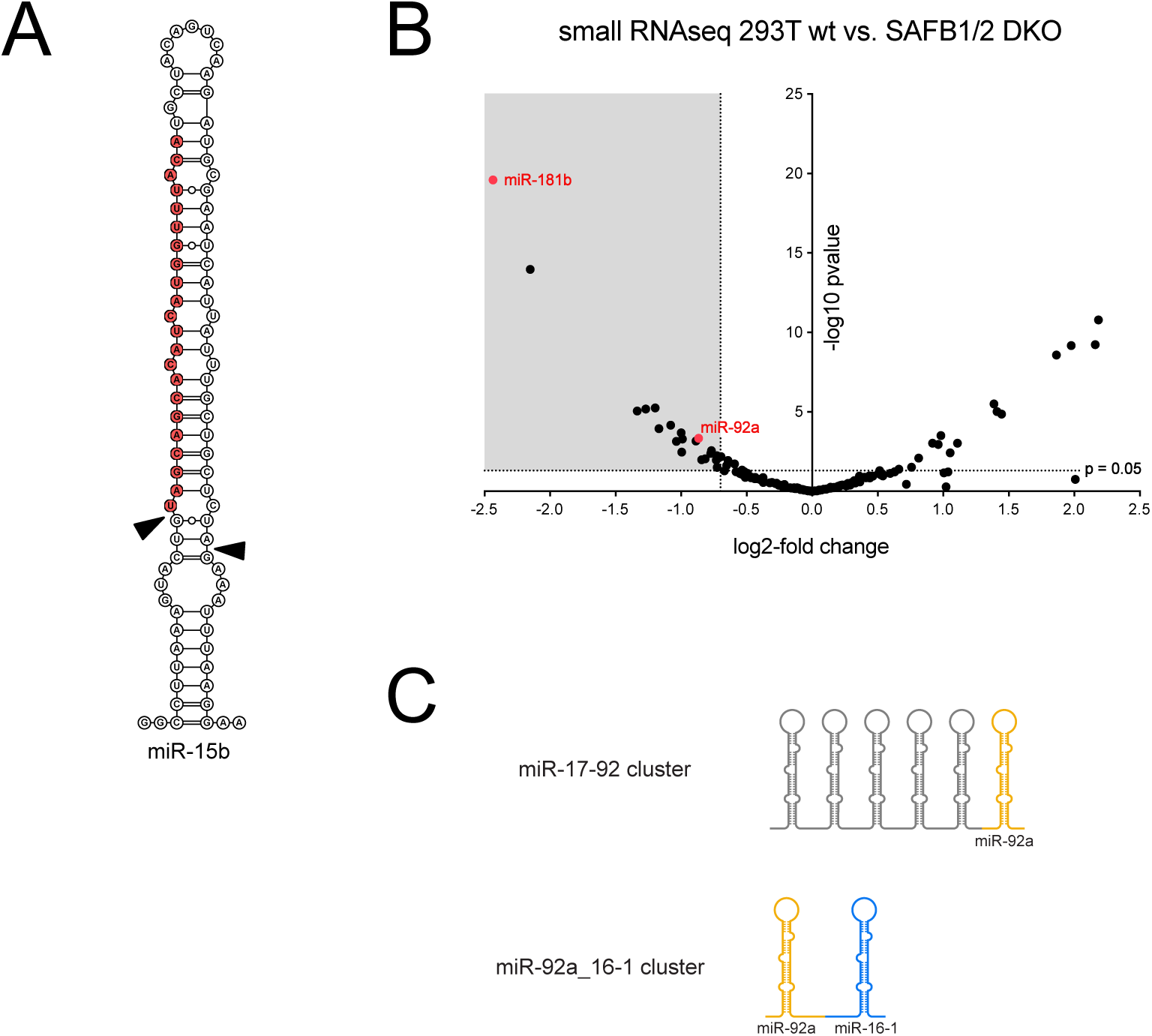
**A**. Schematic illustration of the stem-loop portion of primary miR-15b. Red nucleotides correspond to the respective mature miRNA. Microprocessor cleavage sites are marked by black triangles. **B**. Small RNA sequencing identifies several miRNAs that are reduced upon SAFB1/2 deletion in 293T cells. Total RNA derived from Safb1/2 DKO 293T cells and the corresponding controls was analyzed for global miRNA expression by small RNA sequencing. The grey square marks miRNAs that are significantly reduced in the absence of SAFB1 and 2 (reduction more than 0,7-fold (log2); p < 0.05). miR-181b and miR-92a chosen for further analysis are depicted in red. **C**. Schematic illustration of the chimeric miR-92a_16-1 cluster. To investigate a putative dependency of miR-92a on cluster assistance, it was combined with miR-16-1 to be expressed as a clustered primary miRNA.

**Figure S5.**
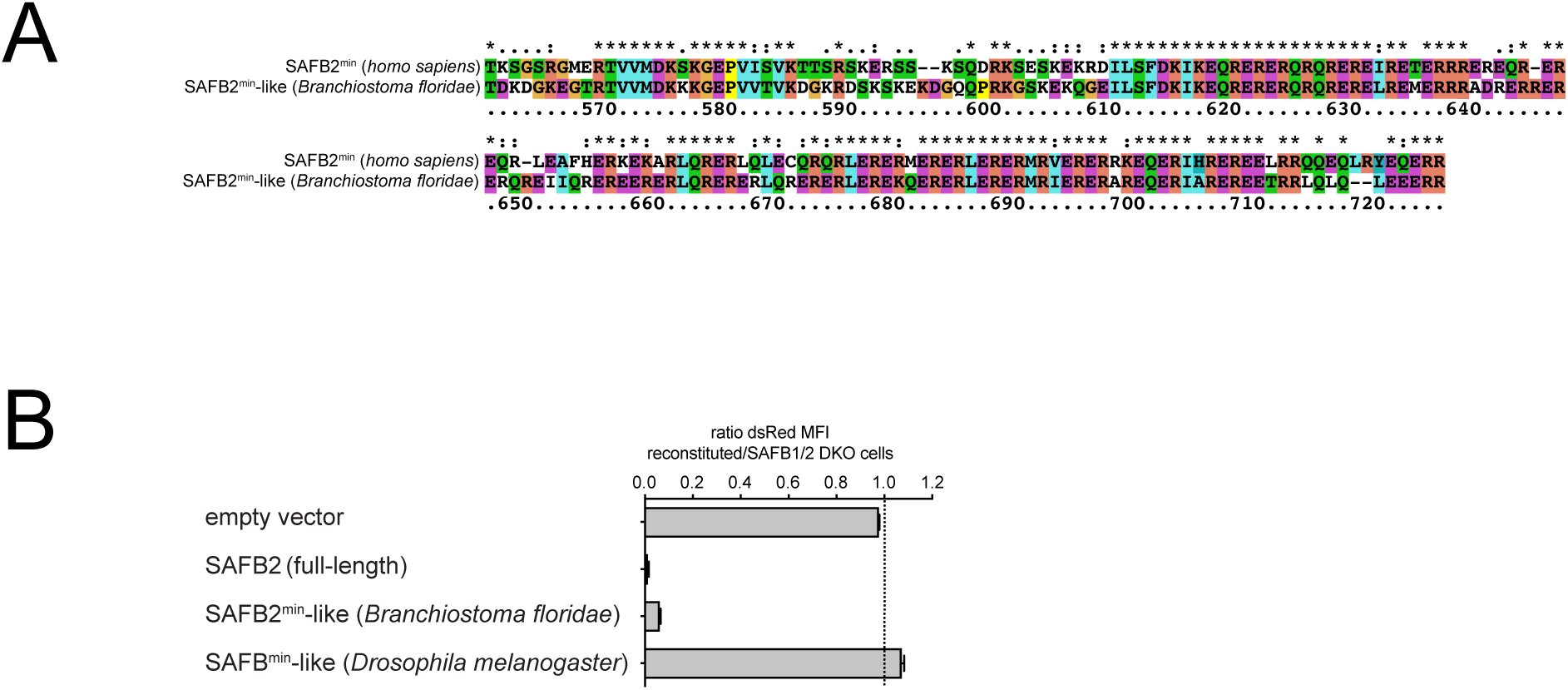
**A**. Alignment of the human SAFB2^min^ fragment and the corresponding SAFB2-like homolog in *Branchiostoma floridae*. Numbers correspond to the respective aa position in the human protein. **B**. SAFB2^min^ function is highly conserved among chordates. Screen cells lacking SAFB1 and 2 and sorted for the dsRed+ population were reconstituted with full-length human SAFB2 or with SAFB2^min^-like variants of *Branchiostoma floridae* and *Drosophila melanogaster* (A and data not shown). MiR-15a^mut^ reporter expression was analyzed by flow cytometry after one week. The individual bars show the ratio of dsRed MFI comparing transduced versus non-transduced cells. A dashed line marks a ratio of 1, which corresponds to no effect.

## References

Afgan, E., Baker, D., Batut, B., van den Beek, M., Bouvier, D., Čech, M., Chilton, J., Clements, D., Coraor, N., Grüning, B.A., Guerler, A., Hillman-Jackson, J., Hiltemann, S., Jalili, V., Rasche, H., Soranzo, N., Goecks, J., Taylor, J., Nekrutenko, A., Blankenberg, D., 2018. The Galaxy platform for accessible, reproducible and collaborative biomedical analyses: 2018 update. Nucleic Acids Res 46, W537–W544. doi:10.1093/nar/gky379

Alarcón, C.R., Lee, H., Goodarzi, H., Halberg, N., Tavazoie, S.F., 2015. N6-methyladenosine marks primary microRNAs for processing. Nature 519, 482–485. doi:10.1038/nature14281

Alfonso-Parra, C., Maggert, K.A., 2010. Drosophila SAF-B links the nuclear matrix, chromosomes, and transcriptional activity. PLoS ONE 5, e10248. doi:10.1371/journal.pone.0010248

Allegra, D., Mertens, D., 2011. In-vivo quantification of primary microRNA processing by Drosha with a luciferase based system. Biochemical and Biophysical Research Communications 406, 501–505. doi:10.1016/j.bbrc.2011.02.055

Altmeyer, M., Toledo, L., Gudjonsson, T., Grøfte, M., Rask, M.-B., Lukas, C., Akimov, V., Blagoev, B., Bartek, J., Lukas, J., 2013. The chromatin scaffold protein SAFB1 renders chromatin permissive for DNA damage signaling. Mol. Cell 52, 206–220. doi:10.1016/j.molcel.2013.08.025

Auyeung, V.C., Ulitsky, I., McGeary, S.E., Bartel, D.P., 2013. Beyond secondary structure: primary-sequence determinants license pri-miRNA hairpins for processing. Cell 152, 844–858. doi:10.1016/j.cell.2013.01.031

Baltz, A.G., Munschauer, M., Schwanhäusser, B., Vasile, A., Murakawa, Y., Schueler, M., Youngs, N., Penfold-Brown, D., Drew, K., Milek, M., Wyler, E., Bonneau, R., Selbach, M., Dieterich, C., Landthaler, M., 2012. The mRNA-bound proteome and its global occupancy profile on protein-coding transcripts. Mol. Cell 46, 674–690. doi:10.1016/j.molcel.2012.05.021

Castello, A., Fischer, B., Eichelbaum, K., Horos, R., Beckmann, B.M., Strein, C., Davey, N.E., Humphreys, D.T., Preiss, T., Steinmetz, L.M., Krijgsveld, J., Hentze, M.W., 2012. Insights into RNA Biology from an Atlas of Mammalian mRNA-Binding Proteins. Cell. doi:10.1016/j.cell.2012.04.031

Cheng, T.-L., Wang, Z., Liao, Q., Zhu, Y., Zhou, W.-H., Xu, W., Qiu, Z., 2014. MeCP2 suppresses nuclear microRNA processing and dendritic growth by regulating the DGCR8/Drosha complex. Developmental Cell 28, 547–560. doi:10.1016/j.devcel.2014.01.032

Chiodi, I., Biggiogera, M., Denegri, M., Corioni, M., Weighardt, F., Cobianchi, F., Riva, S., Biamonti, G., 2000. Structure and dynamics of hnRNP-labelled nuclear bodies induced by stress treatments. J. Cell. Sci. 113 (Pt 22), 4043–4053.

Dai, Z., Sheridan, J.M., Gearing, L.J., Moore, D.L., Su, S., Wormald, S., Wilcox, S., O’Connor, L., Dickins, R.A., Blewitt, M.E., Ritchie, M.E., 2014. edgeR: a versatile tool for the analysis of shRNA-seq and CRISPR-Cas9 genetic screens. F1000Res 3, 95. doi:10.12688/f1000research.3928.2

Darty, K., Denise, A., Ponty, Y., 2009. VARNA: Interactive drawing and editing of the RNA secondary structure. Bioinformatics 25, 1974–1975. doi:10.1093/bioinformatics/btp250

Denegri, M., Chiodi, I., Corioni, M., Cobianchi, F., Riva, S., Biamonti, G., 2001. Stress-induced nuclear bodies are sites of accumulation of pre-mRNA processing factors. Mol. Biol. Cell 12, 3502–3514. doi:10.1091/mbc.12.11.3502

Denli, A.M., Tops, B.B.J., Plasterk, R.H.A., Ketting, R.F., Hannon, G.J., 2004. Processing of primary microRNAs by the Microprocessor complex. Nature 432, 231–235. doi:10.1038/nature03049

Djuranovic, S., Nahvi, A., Green, R., 2012. miRNA-mediated gene silencing by translational repression followed by mRNA deadenylation and decay. Science 336, 237–240. doi:10.1126/science.1215691

Fang, W., Bartel, D.P., 2015. The Menu of Features that Define Primary MicroRNAs and Enable De Novo Design of MicroRNA Genes. Mol. Cell 60, 131–145. doi:10.1016/j.molcel.2015.08.015

Fletcher, C.E., Godfrey, J.D., Shibakawa, A., Bushell, M., Bevan, C.L., 2016. A novel role for GSK3β as a modulator of Drosha microprocessor activity and MicroRNA biogenesis. Nucleic Acids Res. doi:10.1093/nar/gkw938

Friedman, R.C., Farh, K.K.H., Burge, C.B., Bartel, D.P., 2008. Most mammalian mRNAs are conserved targets of microRNAs. Genome Res. 19, 92–105. doi:10.1101/gr.082701.108

Gregory, R.I., Yan, K.-P., Amuthan, G., Chendrimada, T., Doratotaj, B., Cooch, N., Shiekhattar, R., 2004. The Microprocessor complex mediates the genesis of microRNAs. Nature 432, 235–240. doi:10.1038/nature03120

Grishok, A., Pasquinelli, A.E., Conte, D., Li, N., Parrish, S., Ha, I., Baillie, D.L., Fire, A., Ruvkun, G., Mello, C.C., 2001. Genes and mechanisms related to RNA interference regulate expression of the small temporal RNAs that control C. elegans developmental timing. Cell 106, 23–34. doi:10.1016/s0092-8674(01)00431-7

Guo, H., Ingolia, N.T., Weissman, J.S., Bartel, D.P., 2010. Mammalian microRNAs predominantly act to decrease target mRNA levels. Nature 466, 835–840. doi:10.1038/nature09267

Haar, J., Contrant, M., Bernhardt, K., Feederle, R., Diederichs, S., Pfeffer, S., Delecluse, H.- J., 2016. The expression of a viral microRNA is regulated by clustering to allow optimal B cell transformation. Nucleic Acids Res 44, 1326–1341. doi:10.1093/nar/gkv1330

Hafner, M., Landthaler, M., Burger, L., Khorshid, M., Hausser, J., Berninger, P., Rothballer, A., Ascano, M., Jungkamp, A.-C., Munschauer, M., Ulrich, A., Wardle, G.S., Dewell, S., Zavolan, M., Tuschl, T., 2010. Transcriptome-wide identification of RNA-binding protein and microRNA target sites by PAR-CLIP. Cell 141, 129–141. doi:10.1016/j.cell.2010.03.009

Han, J., Lee, Y., Yeom, K.-H., Kim, Y.-K., Jin, H., Kim, V.N., 2004. The Drosha-DGCR8 complex in primary microRNA processing. Genes & Development 18, 3016–3027. doi:10.1101/gad.1262504

Han, J., Lee, Y., Yeom, K.-H., Nam, J.-W., Heo, I., Rhee, J.-K., Sohn, S.Y., Cho, Y., Zhang, B.-T., Kim, V.N., 2006. Molecular basis for the recognition of primary microRNAs by the Drosha-DGCR8 complex. Cell 125, 887–901. doi:10.1016/j.cell.2006.03.043

Hernández-Hernández, J.M., Mallappa, C., Nasipak, B.T., Oesterreich, S., Imbalzano, A.N., 2013. The Scaffold attachment factor b1 (Safb1) regulates myogenic differentiation by facilitating the transition of myogenic gene chromatin from a repressed to an activated state. Nucleic Acids Res 41, 5704–5716. doi:10.1093/nar/gkt285

Hutvágner, G., McLachlan, J., Pasquinelli, A.E., Bálint, E., Tuschl, T., Zamore, P.D., 2001. A cellular function for the RNA-interference enzyme Dicer in the maturation of the let-7 small temporal RNA. Science 293, 834–838. doi:10.1126/science.1062961

Ivanova, M., Dobrzycka, K.M., Jiang, S., Michaelis, K., Meyer, R., Kang, K., Adkins, B., Barski, O.A., Zubairy, S., Divisova, J., Lee, A.V., Oesterreich, S., 2005. Scaffold attachment factor B1 functions in development, growth, and reproduction. Mol. Cell. Biol. 25, 2995–3006. doi:10.1128/MCB.25.8.2995-3006.2005

Jiang, S., Katz, T.A., Garee, J.P., DeMayo, F.J., Lee, A.V., Oesterreich, S., 2015. Scaffold attachment factor B2 (SAFB2)-null mice reveal non-redundant functions of SAFB2 compared with its paralog, SAFB1. Dis Model Mech 8, 1121–1127. doi:10.1242/dmm.019885

Ketting, R.F., Fischer, S.E., Bernstein, E., Sijen, T., Hannon, G.J., Plasterk, R.H., 2001. Dicer functions in RNA interference and in synthesis of small RNA involved in developmental timing in C. elegans. Genes & Development 15, 2654–2659. doi:10.1101/gad.927801

Kiledjian, M., Dreyfuss, G., 1992. Primary structure and binding activity of the hnRNP U protein: binding RNA through RGG box. EMBO J. 11, 2655–2664.

Kim, D., Langmead, B., Salzberg, S.L., 2015. HISAT: a fast spliced aligner with low memory requirements. Nature Methods 12, 357–360. doi:10.1038/nmeth.3317

Knackmuss, U., Lindner, S.E., Aneichyk, T., Kotkamp, B., Knust, Z., Villunger, A., Herzog, S., 2015. MAP3K11 is a tumor suppressor targeted by the oncomiR miR-125b in early B cells. Cell Death Differ. doi:10.1038/cdd.2015.87

Kozomara, A., Griffiths-Jones, S., 2011. miRBase: integrating microRNA annotation and deep-sequencing data. Nucleic Acids Res 39, D152–7. doi:10.1093/nar/gkq1027

Kwon, S.C., Nguyen, T.A., Choi, Y.-G., Jo, M.H., Hohng, S., Kim, V.N., Woo, J.-S., 2016. Structure of Human DROSHA. Cell 164, 81–90. doi:10.1016/j.cell.2015.12.019

Landthaler, M., Yalcin, A., Tuschl, T., 2004. The human DiGeorge syndrome critical region gene 8 and Its D. melanogaster homolog are required for miRNA biogenesis. Curr. Biol. 14, 2162–2167. doi:10.1016/j.cub.2004.11.001

Langmead, B., Salzberg, S.L., 2012. Fast gapped-read alignment with Bowtie 2. Nature Methods 9, 357–359. doi:10.1038/nmeth.1923

Lataniotis, L., Albrecht, A., Kok, F.O., Monfries, C.A.L., Benedetti, L., Lawson, N.D., Hughes, S.M., Steinhofel, K., Mayr, M., Zampetaki, A., 2017. CRISPR/Cas9 editing reveals novel mechanisms of clustered microRNA regulation and function. Sci Rep 7, 8585. doi:10.1038/s41598-017-09268-0

Lee, Y., Ahn, C., Han, J., Choi, H., Kim, J., Yim, J., Lee, J., Provost, P., Rådmark, O., Kim, S., Kim, V.N., 2003. The nuclear RNase III Drosha initiates microRNA processing. Nature 425, 415–419. doi:10.1038/nature01957

Lim, L.P., Glasner, M.E., Yekta, S., Burge, C.B., Bartel, D.P., 2003. Vertebrate microRNA genes. Science 299, 1540. doi:10.1126/science.1080372

Liu, H.-W., Banerjee, T., Guan, X., Freitas, M.A., Parvin, J.D., 2015. The chromatin scaffold protein SAFB1 localizes SUMO-1 to the promoters of ribosomal protein genes to facilitate transcription initiation and splicing. Nucleic Acids Res 43, 3605–3613. doi:10.1093/nar/gkv246

Liu, J., Carmell, M.A., Rivas, F.V., Marsden, C.G., Thomson, J.M., Song, J.-J., Hammond, S.M., Joshua-Tor, L., Hannon, G.J., 2004. Argonaute2 is the catalytic engine of mammalian RNAi. Science 305, 1437–1441. doi:10.1126/science.1102513

Liu, R., Holik, A.Z., Su, S., Jansz, N., Chen, K., Leong, H.S., Blewitt, M.E., Asselin-Labat, M.-L., Smyth, G.K., Ritchie, M.E., 2015. Why weight? Modelling sample and observational level variability improves power in RNA-seq analyses. Nucleic Acids Res 43, e97. doi:10.1093/nar/gkv412

Livak, K.J., Schmittgen, T.D., 2001. Analysis of relative gene expression data using real-time quantitative PCR and the 2(-Delta Delta C(T)) Method. Methods 25, 402–408. doi:10.1006/meth.2001.1262

Lorenz, R., Bernhart, S.H., Höner Zu Siederdissen, C., Tafer, H., Flamm, C., Stadler, P.F., Hofacker, I.L., 2011. ViennaRNA Package 2.0. Algorithms Mol Biol 6, 26. doi:10.1186/1748-7188-6-26

Madeira, F., Park, Y.M., Lee, J., Buso, N., Gur, T., Madhusoodanan, N., Basutkar, P., Tivey, A.R.N., Potter, S.C., Finn, R.D., Lopez, R., 2019. The EMBL-EBI search and sequence analysis tools APIs in 2019. Nucleic Acids Res 47, W636–W641. doi:10.1093/nar/gkz268

Nguyen, T.A., Jo, M.H., Choi, Y.-G., Park, J., Kwon, S.C., Hohng, S., Kim, V.N., Woo, J.-S., 2015. Functional Anatomy of the Human Microprocessor. Cell 161, 1374–1387. doi:10.1016/j.cell.2015.05.010

Nussbacher, J.K., Yeo, G.W., 2018. Systematic Discovery of RNA Binding Proteins that Regulate MicroRNA Levels. Mol. Cell 69, 1005–1016.e7. doi:10.1016/j.molcel.2018.02.012

Oesterreich, S., Lee, A.V., Sullivan, T.M., Samuel, S.K., Davie, J.R., Fuqua, S.A., 1997. Novel nuclear matrix protein HET binds to and influences activity of the HSP27 promoter in human breast cancer cells. J. Cell. Biochem. 67, 275–286.

Omura, Y., Nishio, Y., Takemoto, T., Ikeuchi, C., Sekine, O., Morino, K., Maeno, Y., Obata, T., Ugi, S., Maegawa, H., Kimura, H., Kashiwagi, A., 2009. SAFB1, an RBMX-binding protein, is a newly identified regulator of hepatic SREBP-1c gene. BMB Rep 42, 232– 237.

Park, J.W., Parisky, K., Celotto, A.M., Reenan, R.A., Graveley, B.R., 2004. Identification of alternative splicing regulators by RNA interference in Drosophila. Proc. Natl. Acad. Sci. U.S.A. 101, 15974–15979. doi:10.1073/pnas.0407004101

Peidis, P., Voukkalis, N., Aggelidou, E., Georgatsou, E., Hadzopoulou-Cladaras, M., Scott, R.E., Nikolakaki, E., Giannakouros, T., 2011. SAFB1 interacts with and suppresses the transcriptional activity of p53. FEBS Lett. 585, 78–84. doi:10.1016/j.febslet.2010.11.054

Ramanathan, M., Majzoub, K., Rao, D.S., Neela, P.H., Zarnegar, B.J., Mondal, S., Roth, J.G., Gai, H., Kovalski, J.R., Siprashvili, Z., Palmer, T.D., Carette, J.E., Khavari, P.A., 2018. RNA-protein interaction detection in living cells. Nature Methods 15, 207–212. doi:10.1038/nmeth.4601

Reich, M., Liefeld, T., Gould, J., Lerner, J., Tamayo, P., Mesirov, J.P., 2006. GenePattern 2.0. Nat Genet 38, 500–501. doi:10.1038/ng0506-500

Renz, A., Fackelmayer, F.O., 1996. Purification and molecular cloning of the scaffold attachment factor B (SAF-B), a novel human nuclear protein that specifically binds to S/MAR-DNA. Nucleic Acids Res 24, 843–849. doi:10.1093/nar/24.5.843

Rivers, C., Idris, J., Scott, H., Rogers, M., Lee, Y.-B., Gaunt, J., Phylactou, L., Curk, T., Campbell, C., Ule, J., Norman, M., Uney, J.B., 2015. iCLIP identifies novel roles for SAFB1 in regulating RNA processing and neuronal function. BMC Biol. 13, 111. doi:10.1186/s12915-015-0220-7

Robinson, M.D., McCarthy, D.J., Smyth, G.K., 2010. edgeR: a Bioconductor package for differential expression analysis of digital gene expression data. Bioinformatics 26, 139–140. doi:10.1093/bioinformatics/btp616

Runte, M., Hüttenhofer, A., Gross, S., Kiefmann, M., Horsthemke, B., Buiting, K., 2001. The IC-SNURF-SNRPN transcript serves as a host for multiple small nucleolar RNA species and as an antisense RNA for UBE3A. Hum. Mol. Genet. 10, 2687–2700. doi:10.1093/hmg/10.23.2687

Sanjana, N.E., Shalem, O., Zhang, F., 2014. Improved vectors and genome-wide libraries for CRISPR screening. Nature Methods 11, 783–784. doi:10.1038/nmeth.3047

Schwarz, D.S., Hutvagner, G., Du, T., Xu, Z., Aronin, N., Zamore, P.D., 2003. Asymmetry in the assembly of the RNAi enzyme complex. Cell 115, 199–208. doi:10.1016/s0092-8674(03)00759-1

Sergeant, K.A., Bourgeois, C.F., Dalgliesh, C., Venables, J.P., Stevenin, J., Elliott, D.J., 2007. Alternative RNA splicing complexes containing the scaffold attachment factor SAFB2. J. Cell. Sci. 120, 309–319. doi:10.1242/jcs.03344

Shalem, O., Sanjana, N.E., Hartenian, E., Shi, X., Scott, D.A., Mikkelson, T., Heckl, D., Ebert, B.L., Root, D.E., Doench, J.G., Zhang, F., 2014. Genome-scale CRISPR-Cas9 knockout screening in human cells. Science 343, 84–87. doi:10.1126/science.1247005

Townson, S.M., Dobrzycka, K.M., Lee, A.V., Air, M., Deng, W., Kang, K., Jiang, S., Kioka, N., Michaelis, K., Oesterreich, S., 2003. SAFB2, a new scaffold attachment factor homolog and estrogen receptor corepressor. J. Biol. Chem. 278, 20059–20068. doi:10.1074/jbc.M212988200

Treiber, T., Treiber, N., Plessmann, U., Harlander, S., Daiß, J.-L., Eichner, N., Lehmann, G., Schall, K., Urlaub, H., Meister, G., 2017. A Compendium of RNA-Binding Proteins that Regulate MicroRNA Biogenesis. Mol. Cell 66, 270–284.e13. doi:10.1016/j.molcel.2017.03.014

Truscott, M., Islam, A.B.M.M.K., Frolov, M.V., 2016. Novel regulation and functional interaction of polycistronic miRNAs. RNA 22, 129–138. doi:10.1261/rna.053264.115

Tu, C.-C., Zhong, Y., Nguyen, L., Tsai, A., Sridevi, P., Tarn, W.-Y., Wang, J.Y.J., 2015. The kinase ABL phosphorylates the microprocessor subunit DGCR8 to stimulate primary microRNA processing in response to DNA damage. Sci Signal 8, ra64. doi:10.1126/scisignal.aaa4468

Varkonyi-Gasic, E., Wu, R., Wood, M., Walton, E.F., Hellens, R.P., 2007. Protocol: a highly sensitive RT-PCR method for detection and quantification of microRNAs. Plant Methods 3, 12. doi:10.1186/1746-4811-3-12

Wang, Y., Zhang, X., Zhang, H., Lu, Y., Huang, H., Dong, X., Chen, J., Dong, J., Yang, X., Hang, H., Jiang, T., 2012. Coiled-coil networking shapes cell molecular machinery. Mol. Biol. Cell 23, 3911–3922. doi:10.1091/mbc.E12-05-0396

Yamaguchi, A., Takanashi, K., 2016. FUS interacts with nuclear matrix-associated protein SAFB1 as well as Matrin3 to regulate splicing and ligand-mediated transcription. Sci Rep 6, 35195. doi:10.1038/srep35195

Zeng, Y., Cullen, B.R., 2005. Efficient processing of primary microRNA hairpins by Drosha requires flanking nonstructured RNA sequences. J. Biol. Chem. 280, 27595–27603. doi:10.1074/jbc.M504714200

Zeng, Y., Cullen, B.R., 2003. Sequence requirements for micro RNA processing and function in human cells. RNA 9, 112–123.

Zeng, Y., Yi, R., Cullen, B.R., 2005. Recognition and cleavage of primary microRNA precursors by the nuclear processing enzyme Drosha. EMBO J. 24, 138–148. doi:10.1038/sj.emboj.7600491

